# Rickettsiales’ deep evolutionary history sheds light on the emergence of intracellular lifestyles

**DOI:** 10.1101/2023.01.31.526412

**Authors:** Lucas Serra Moncadas, Tanja Shabarova, Vinicius Silva Kavagutti, Paul-Adrian Bulzu, Maria-Cecilia Chiriac, Soo-Je Park, Indranil Mukherjee, Rohit Ghai, Adrian-Stefan Andrei

## Abstract

Ancient bacteria largely lived and flourished as free-living cells till the rise of eukaryotes triggered their adaptation to a new habitat: the intracellular milieu. Rickettsiales bacteria are the most prevalent intracellular microorganisms discovered and the culprits behind some of mankind’s worst pestilential diseases. Here, we show that intracellularity is not a defining feature of the group and describe the eco-evolutionary processes that transformed harmless free-living bacteria into obligate intracellular symbionts and parasites. We found that the evolution of free-living lineages towards enhanced cross-feeding interactions with microbial eukaryotes trapped them in a nutritional bind to their trophic partners. We discovered that the oldest Rickettsiales lineages are the closest relatives to modern eukaryotes and are enriched in proteins predicted to have been present in the mitochondrial ancestor. This study not only opens avenues for the detection and surveillance of emerging diseases but also expands our understanding of the origins of complex life.

**One-Sentence Summary:** Cross-feeding interactions shaped Rickettsiales genomic architectures along the parasite/free-living spectrum

## Main Text

The emergence of bacterial intracellular lifestyle occupies a relatively recent chapter in the history of life on our planet. This ability, which allowed bacteria to live and thrive within the cells of other organisms, irreversibly changed the course of evolution by enabling the emergence of complex life (through eukaryogenesis) and establishing the foundation for many interspecies symbiotic interactions. The Rickettsiales embody some of the most notorious branches of the bacterial line of descent. Their renowned reputation stems from the accommodation of lineages accountable for some of the deadliest and oldest pestilential diseases of mankind^1^ and their nexus with eukaryogenesis^2^: the emergence of eukaryotes. The order-level division, which draws its name from the causative agent of epidemic typhus *Rickettsia prowazekii*, shepherded humanity throughout the ages as shown by time-worn records extending from antiquity (e.g. Plague of Athens) to modernity (e.g. Burundian civil wars)^1^. Even though the death toll exacted by typhus (in the order of millions) likely surpasses that of military conflicts, its present-day grip on humanity has only been loosened by the onset of the antibiotic era^3^. Mankind’s struggle to subdue one of its most dreaded nemesis generated a wealth of knowledge that permeated profoundly the biological sciences, shaping the fields of epidemiology, immunology, and evolutionary biology. Nowadays, Rickettsiales appear outstripped of their infamy, emerging instead as the utmost abundant clade of obligate intracellular bacteria^4^ that engage in a wide spectrum of trophic interactions (from mutualism to parasitism) with a plethora of eukaryotic hosts^5^.

The early dawn of genomics realization that Rickettsiales’ intracellular lifestyles molded their genomic architectures crystalized later into the mainstream reductive genome evolution paradigm^6^. In light of this, bacterial genomic architecture is governed by genetic drift, which acts towards eroding superfluous genomic regions, particularly in lineages that experience population bottlenecks imposed by host associations^7^. Although intracellularity, genome shrinkage, and impaired metabolic capacity are key features of reductive genome evolution, the ecological pressures and evolutionary trajectories that generated them (in Rickettsiales) have remained obscure. The present work integrates Rickettsiales genomic architecture into an evolutionary framework to reveal the processes that drove environmental free-living cells to live inside of other cells. This was largely made possible by the retrieval of genetic information from diverse and undescribed aquatic lineages.

### Evolutionary history

The mainstream view portraying Rickettsiales as intracellular symbionts of arthropods and vertebrates^8^ has been recently challenged by their detection in environmental diversity surveys^9,10^ and the discovery of lineages associated with aquatic unicellular eukaryotes^11–13^. As genome-to-habitat matchings of publicly available Rickettsiales genomes indicated novel diversity within freshwater environments (GTDB R05-RS95 database; see Supplementary Data S1), the data-generating strategy was focused on likewise habitats (see Supplementary Text). A database of approximately 6 000 curated metagenome-assembled genomes (MAGs) was constructed by employing genome-resolved metagenomics techniques on obtained environmental shotgun sequencing data. Phylogenomics-based taxonomic classification identified 49 MAGs as belonging to the order Rickettsiales (Fig. S1). This dataset was expanded with the addition of all representative genomes/MAGs (n=80) available in the public repositories (GTDB R05-RS95 database). Genomic-recovered 16S rRNA sequences used to anchor the Rickettsiales dataset (n=129 MAGs/genomes) into the larger gene-based bacterial taxonomy showed i) recovery of novel family-level clades and ii) existence of lineages (i.e. within family UBA1997) that likely evaded diversity surveys owing to classification within the mitochondrial line of descent (see Supplementary Text; Fig. S2). As family-level categories likely encompass congruent ecological strategies^14^, they were used as scaffolds for coupling phylogeny with metabolic pathway radiation. The depicted evolutionary history reconstruction (Fig. 1) was selected based on goodness-of-fit evaluations of three posterior consensus phylogenies produced under different combinations of empirical profiles and infinite mixture models (Figs. S3, S4). The mapping of genome-level metabolic reconstructions on the tree’s topological backbone brought to the fore the division of the Rickettsiales’ evolutionary history into three major stages: i) intracellular, ii) transition and iii) ancient.

**Fig. 1.**
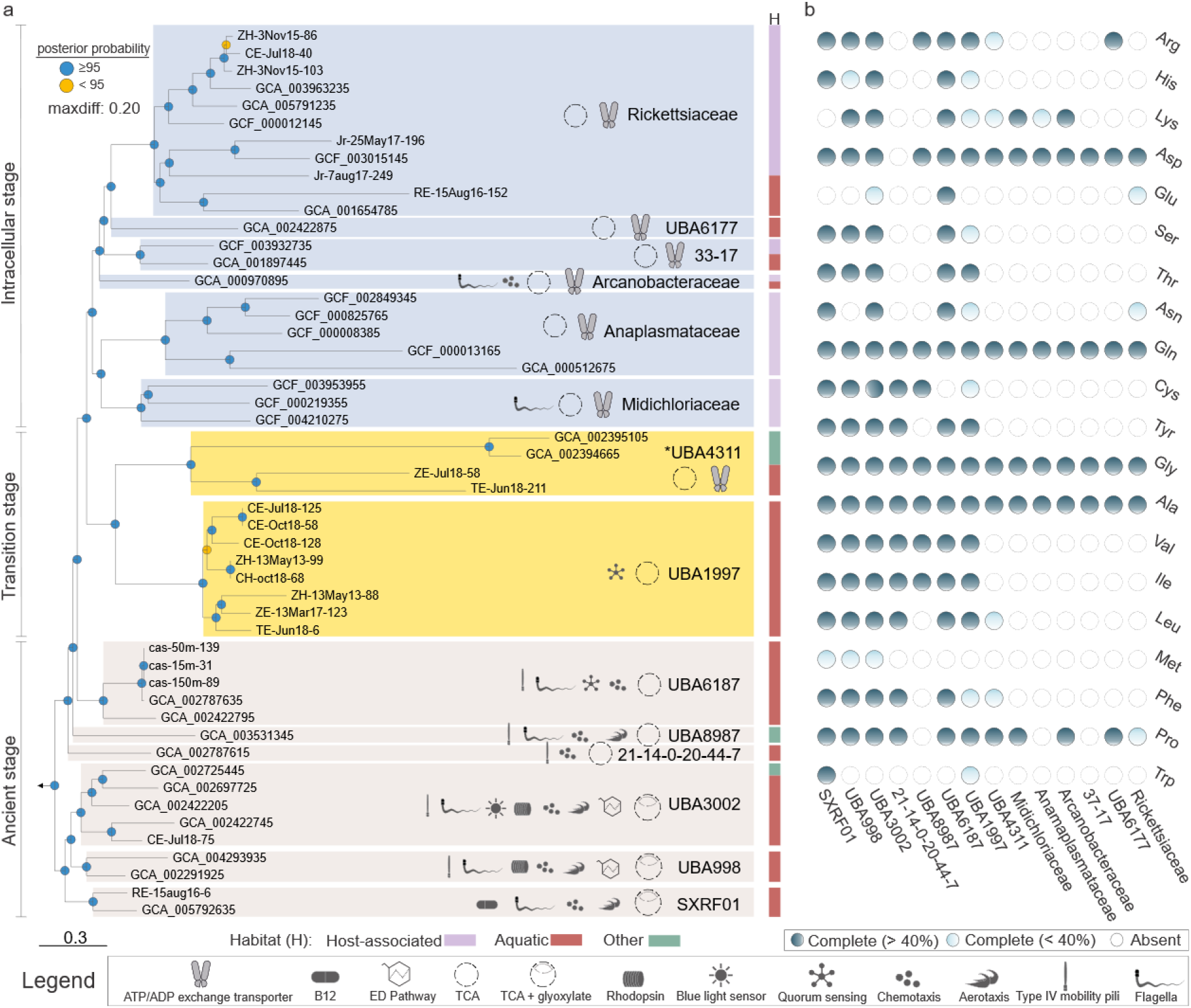
Rickettsiales evogenomic investigations. a) Family-centric genome-based phylogeny generated through Bayesian inference (-cat -f81 -dgam4). Posterior probability values are represented through colored circles (top left legend). The scale bar indicates the number of substitutions per site. The habitat of origin (H) is shown on the right side of the panel (bottom left legend). The pictograms reflect family-level physiological features obtained through genome-informed metabolic reconstructions (bottom panel inset). b) Census of metabolic pathways responsible for proteinogenic amino acid biosynthesis. The legend (bottom right) specifies the percentage (at the family level) of genomes with complete metabolic pathways. ^*^Taxonomic clade (UBA4311) situated at the lower limit of family-level similarity.

The intracellular stage (resolved with a posterior probability of 0.95-1) depicts the latest diversification of the order and encompasses the iconic symbiotic families of radiation (Rickettsiales *sensu stricto*). Most of the six lineages (Midichloraceae to Rickettsiaceae) are host-associated or found in aquatic habitats where they likely engaged in symbiosis with microbial eukaryotes^15^, as this monophyletic stage was marked by severe impairment in amino acid biosynthetic capacity and the presence of ATP/ADP exchange transporter (Fig. 1). This unusual translocase represents a hallmark of intracellular lifestyle in Rickettsiales, as it allows bacteria to perform ‘energy parasitism’ by importing host-derived energy-rich ATP in exchange for endogenous produced ADP. As the adaptation to the intracellular milieu increased, the ATP/ADP translocase paralogues expanded functionality in the youngest lineages of the stage (i.e. UBA6177/Rickettsiaceae branch; Fig. S5) through increasing imported nucleotides diversity.

The transition stage (resolved with posterior probability 1) is portrayed by two families (UBA1997 and UBA4311) that share common ancestry with the intracellular one with whom they are connected through a sister-clade relationship (Fig. 1). Though phylogenetically linked, these two families exhibit distinct physiological characteristics. While UBA4311 lineage is typified by a lack of amino acid biosynthetic pathways and likely hosts the earliest intracellular representatives (as indicated by the presence of ATP/ADP translocase), UBA1997 differentiates through substantial augmentation of amino acid biosynthetic machinery and capacity for gene expression regulation through quorum signaling. Although UBA4311 encompassed fast-evolving lineages within the lower limit of family-level similarity found in aquatic and gut microbiomes, UBA1997 seemed to be restricted to freshwater ecosystems.

The ancient stage (resolved with posterior probability 0.99-1) encompasses six basal families (SXRF01 to UBA6187) that collectively comprise the oldest evolutionary lineages within Rickettsiales. This stage comprises taxa generally present in aquatic habitats that are evolutionarily linked and physiologically typified by the capacity to detect (i.e. aerotaxis, chemotaxis, blue light sensors, rhodopsins, quorum sensing) and respond through directional motility (i.e., flagella, type IV motility pili) to external environmental stimuli (Fig. 1; Supplementary Text; Fig. S6). An inventory of complete metabolic pathways involved in the biosynthesis of 20 proteinogenic amino acids indicated metabolic ‘autonomy’ since families of the ancient stage showed the capacity to produce most of the required proteome’s building blocks (Fig. 1).

Genome-scale metabolic reconstructions revealed the potentiality for autonomous heterotrophic metabolism in transition and ancient stages. Typified by reduced diversity of organic carbon uptake transporters, stage representatives likely fuel their metabolic circuitry with energy generated through amino acid breakdown (generally L-alanine and/or L-serine). Briefly, imported amino acids (L-type amino acid transporters, Na^+^/alanine symporters) can be degraded to pyruvate (i.e., D-amino acid dehydrogenase, serine deaminase), which could be decarboxylated (pyruvate dehydrogenase complex) to the hub metabolite acetyl-CoA. Through this, amino acid-derived pyruvate can either enter the tricarboxylic acid (TCA) cycle and power aerobic respiration or act as a foundation for macromolecular components biosynthesis (e.g. carbohydrates produced by the glyoxylate cycle). It is noteworthy to mention that basal families of the ancient stage showed versatility in acetyl-CoA pool regeneration, as they displayed the capacity for carbohydrates or fatty acids degradation. In UBA998 imported maltose (malt maltose/maltooligosaccharide transporter) can be hydrolyzed to D-glucose (malZ alpha-glucosidase), phosphorylated (glucokinases), and subsequently degraded to pyruvate through Entner-Doudoroff pathway. At the same time, in SXRF01 the acetyl-CoA pool can be replenished by the breakdown of exogenous fatty acids (through β-oxidation) that entered cells by free diffusion or that were previously membrane-incorporated (fused 2-acylglycerophospho-ethanolamine acyltransferase/acyl-acyl carrier protein synthetase).

### Genomic architectures of free-living lifestyles

Genomic property screenings depicted an intracellular stage characterized by features typifying reductive genome evolution^6^: i) small genomes, ii) reduced coding potential, iii) low GC content, and iv) limited signal transduction and gene regulation capacity (Fig. 2). Similar attributes were found in the transition stage, which differentiated from the intracellular one through an even further decrease in signal transduction capacity. The ancient stage displayed contrasting genomic properties with expansions in all up-mentioned features and bore little similarity to hallmark characteristics of reductive evolution (Fig. 2). Here, substantial expansions of signal transduction domains and sigma factors point towards the ability to modulate gene expression in unstable niches, while an increased number of GH16 glycoside hydrolases (i.e. β-glycanases) implied the ability for utilization of phytoplankton-derived polysaccharides^16^.

**Fig. 2.**
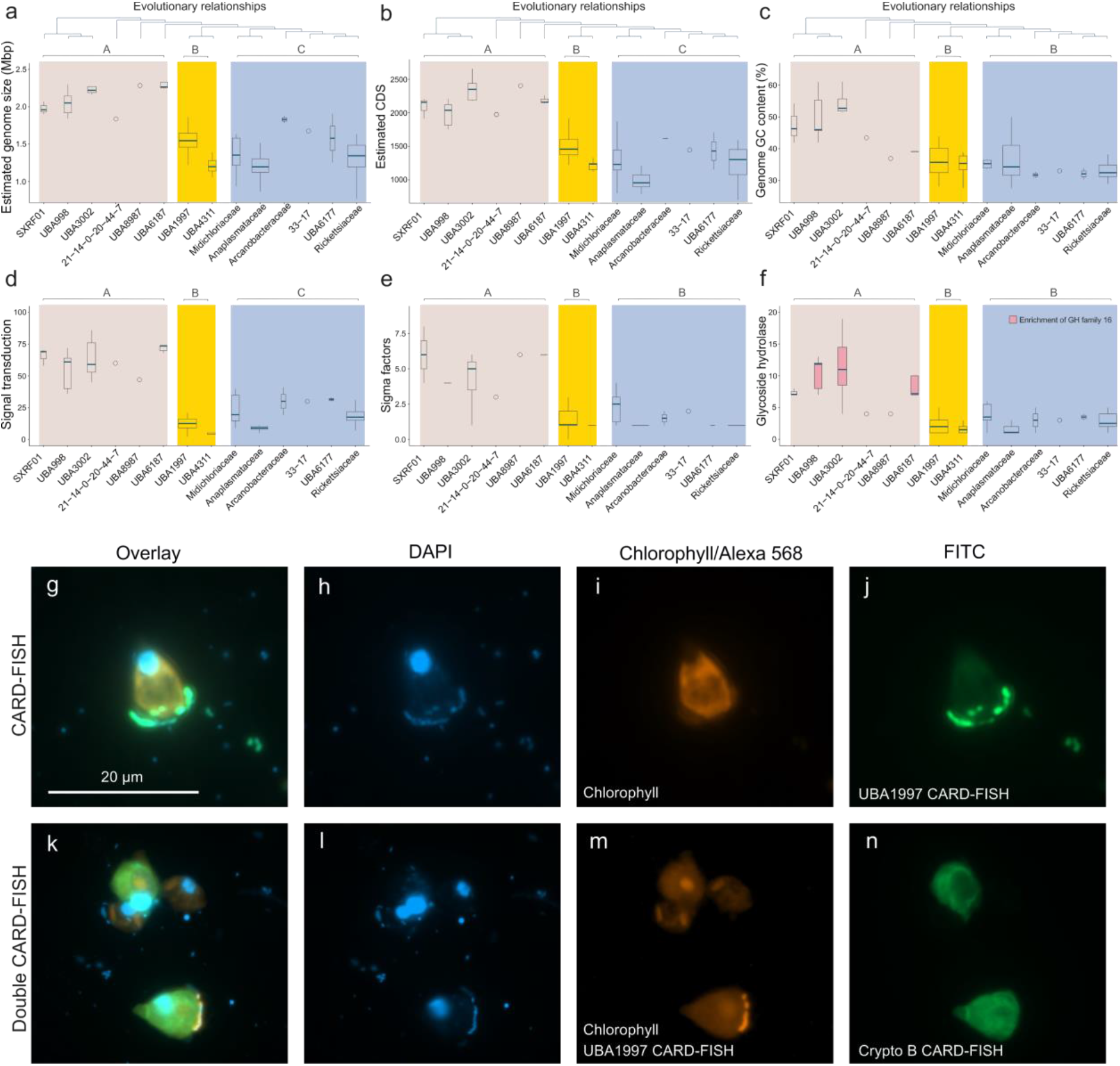
Rickettsiales ecogenomic investigations. a-f) Breakdown of family-level genomic properties and features ordered in a phylogenetic fashion. Distinct evolutionary stages are represented through colored backgrounds. Stage differences are depicted through the usage of uppercase letters (A-C). g-j) Environmental-derived microphotographs displaying UBA1997 family members in epibiotic associations with freshwater microeukaryotes (h-DNA DAPI staining in blue, i-chlorophyll fluorescence in red, j-Catalyzed Reporter Deposition-Fluorescence in situ Hybridization [CARD-FISH] of UBA1997 bacterial ribosomes in green). k-n) Environmental-derived microphotographs linking UBA1997 family members to freshwater cryptophyte algae (l-DNA DAPI staining in blue, m-chlorophyll fluorescence in red, and CARD-FISH of UBA1997 bacterial ribosomes in bright red, n-CARD-FISH of cryptophyte ribosomes in green). The scale bar size corresponds to 20 μm.

Evolutionary history, metabolic capacity, genomic properties, and environmental abundance were further evaluated towards defining suitable targets for phylogenetic staining (through Catalyzed Reporter Deposition-Fluorescence in situ Hybridization, i.e. CARD-FISH). The exclusion of the intracellular stage was motivated by typically low numbers that fell beneath the detection limit (for CARD-FISH in environmental samples) and by the accessibility of ecological data for several members of the clade^8^. UBA1997 was selected as a transition stage representative, as it was the most abundant environmental lineage and presented features that made it comparable to both intracellular (e.g., genomic properties) and ancient (e.g., amino acid metabolism) stages. The environmental presence of the family was assessed by employing designed rRNA-targeting oligonucleotide probes on time series freshwater samples (approx. 225 samples). Microscopic evaluation depicted UBA1997 members as small (length 0.65±0.17 μm, width 0.44±0.09 μm) rod-shaped phenotypes that established persistent surface associations with planktonic unicellular eukaryotes. The identity of the eukaryotic partner was further narrowed down to cryptophyte algae by the usage of rRNA double staining (Fig. 2k-n). Time series profiles depicted UBA1997 as achieving a late spring abundance peak (up to 4% in the surface waters) followed by a sharp decline to a low abundance baseline (0-0.1%) that was maintained throughout the summer/autumn seasons (Supplementary Data S1). Though UBA1997 dynamics seem to mirror the cryptophyte spring bloom event, no further information could be extracted regarding the nature of this epibiotic symbiosis. As a UBA4311 family member (transition stage) was found to engage in extracellular parasitism with aquatic ciliates^13^, it becomes conceivable that UBA1997 representatives could play a role in the environmental food web through modulating cryptophytes’ fitness.

Though all ancient stage families exhibited very low abundances, UBA3002 was targeted for phylogenetic staining, as it comprised a basal branch of the clade and presented genomic and metabolic properties usually associated with free-living lifestyles. The evaluation of the freshwater time series with rRNA-targeting oligonucleotide probes showed the environmental presence of small (length 0.64±0.16 μm, width 0.43±0.07 μm) elongated cocci that appeared either planktonic or scarcely distributed on the surface of Fragilaria diatoms (Fig. S7). The lineage was detected at low abundance (0.1-1.3%) mostly in the surface waters during the summer months (Supplementary Data S1).

### The advent of parasitism

Rickettsiales *sensu stricto* (i.e. intracellular bacteria) are skillful puppeteers that steer the host metabolic machinery toward self-proliferation. Host resource hijacking is usually modulated through moonlighting protein-containing motifs tuned for intramolecular interactions^4^. Among these the most evocative are the ankyrin (ANK) repeat domain-containing proteins, which Rickettsiales maneuver to manipulate a plethora of host cell processes ranging from gene transcription to vesicular trafficking and apoptosis^8^. While previous studies pictured obligate symbiosis as a predictor of ANK repeat abundance^17^ and associated Rickettsiales species^18^ with high ANK numbers, the emergence, and proliferation of these repeats throughout the evolutionary history of the clade have remained unclear.

Evaluation of ANK repeats dynamics within Rickettsiales phylogeny was achieved through juxtaposition with the functionally analogous tetratricopeptide (TPR) ones. The choice stems from functional similarity and distributional ubiquity of TPR across the tree of life^19^ (as ANK exhibits a eukaryotic bias^17^). Genome-centric baseline estimations (using 31 910 representative prokaryotic genomes) highlighted a higher abundance (genome median=10) and spread (detection in 95.1% of genomes) of TPR repeat domain-encoding genes. Disparately, the ANK ones exhibited a narrower distribution (detection in 48.4% of genomes) with lower abundances (genome median=2). The ratio of TPR and ANK repeat domain-encoding genes displayed a dichotomic pattern that mirrored the evolutionary history of Rickettsiales (Fig. 3; Fig. S8). Thus, while the ancient stage families showed abundances comparable with prokaryotic baseline estimations, the transition and intracellular stages displayed a reverse picture with genomic enrichments in ANK- and reductions for TPR-encoding genes (Fig. 3). By connecting the observed patterns of protein scaffold preference with evolutionary history and lifestyle inferences (Fig. 1), it becomes apparent that ANK scaffold selectivity is a late-acquired trait favored by host-associated lineages.

**Fig. 3.**
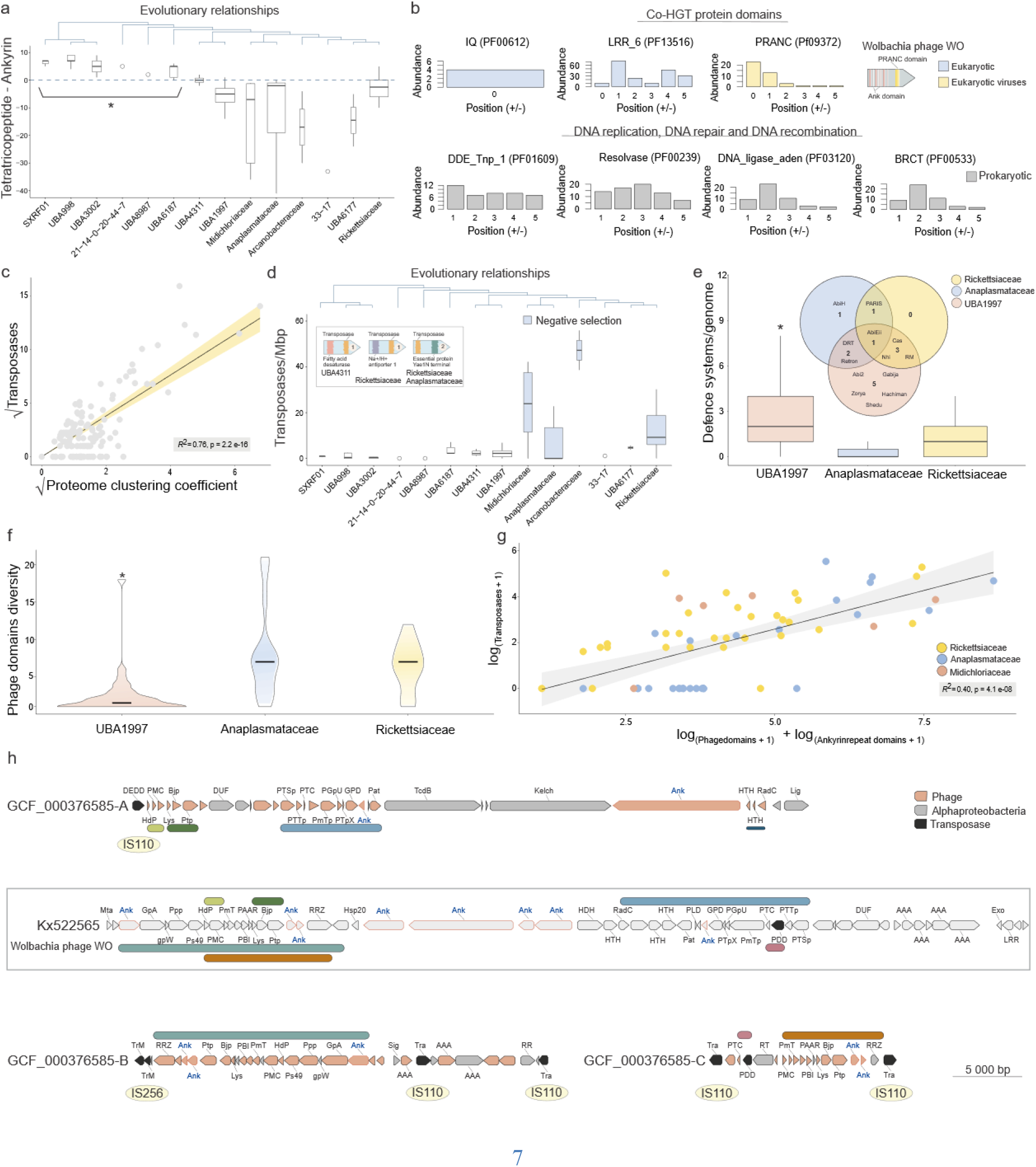
Eco-evolutionary dynamics of ankyrin (ANK) repeat protein domains in Rickettsiales. a) Dynamics of protein-protein interaction domains along Rickettsiales evolutionary history. The panel highlights the emergence of ANK scaffold preference in transition and intracellular stages (^*^statistically significant difference between the ancient stage and the others). b) Ankyrin gene-context analysis showing the abundance of selected co-occurring protein domains (± 5 proteins situated up and downstream of an ANK-containing protein). The colors highlight the inferred evolutionary origin (blue-eukaryotic, yellow-eukaryotic viruses, and grey-bacterial), while the top right inset depicts the schematic representation of a bacteriophage protein (Wolbachia phage WO Kx522565) containing both ANK and PRANC domains. c) Linear regression between the number of transposases and gene duplications (inferred through proteome clustering coefficient) in Rickettsiales genomes. d) Family-level transposase density (transposases/Mbp) depicted in evolutionary order. The colored boxes highlight the selective force (i.e., negative/purifying) governing transposase evolution in major Rickettsiales intracellular lineages. The figure inset depicts the instances (n=4; 0.2% of analyzed transposases) in which transposases were found inserted within metabolic genes. e) Phage defense systems abundance and diversity in transition (UBA1997; n=32) and intracellular (Anaplasmataceae, n=28; Rickettsiaceae, n=30) stages (^*^statistical difference). f) Diversity of phage-derived proteins found in transition (UBA1997; n=32) and intracellular (Anaplasmataceae, n=28; Rickettsiaceae, n=30) stages (^*^statistical difference). g) Multiple regression model between transposase, phage, and ankyrin protein domains. h) Phage-derived genomic regions present in Wolbachia wNo genome (Anaplasmataceae). Synteny blocks are highlighted through color rectangles. The middle inset (grey outline) depicts the Wolbachia WO phage. Gene colors in Wolbachia wNo genome illustrate their provenance (phage-light red, alphaproteobacterial-grey). Transposases are colored black. Transposase-associated insertion sequence types (i.e., IS110 and IS256) are highlighted by yellow backgrounds (not to scale). The scale bar represents 5 000 bp.

To shed light on the origin and mechanisms involved in the maintenance and expansion of the ANK-encoding gene pool a gene context analysis was performed on co-occurring protein domains (within ±5 proteins on each side of a protein having at least one ANK repeat). The results pointed towards a frequent spatial association between ANK and domains involved in DNA breakage, joining and transposition (Fig. 3b; Fig. S9), highlighting the mobilome as a key driver of their propagation (as genome expansions in Rickettsiales were largely associated with transposase duplications; Fig. 3c). While ANK repeat abundance seems to be driven by hitchhiking on mobile genetic elements, the same gene-encoded protein domains revealed glimpses of the evolutionary past. Thus, protein domains of likely eukaryotic/poxviral provenance that resided on the same gene with ANK domain were found to be molecular relicts of long-haul horizontal gene transfer (HGT) events that bridged idiosyncratic domains of life (i.e. Rickettsiales, phages, eukaryotes, and their viruses; see Supplementary Text) (Fig. 3a; Fig. S10). The dispersion of ANK-encoded genes within Rickettsiales lineages was found to be facilitated by temperate phages, as such domains were identified in both prophages and phage genomes (Fig. 3h). Moreover, direct correlations between prophage and ANK encoding genes (Fig. 3g; Fig. S11) point towards the commonality of such HGT events.

Though ANK-carrying phages likely enhance Rickettsiale’s survival in the eukaryotic intracellular milieu^8,20^, their interception by anti-phage defense systems could nullify the adaptive advantage provided by their genomic cargo. The survey of anti-phage defense systems diversity showed a decline in the intracellular ANK-rich lineages (e.g. Anaplasmataceae and Rickettsiaceae), which was corroborated by higher numbers of prophages and prophage remnants (phage-related protein domains diversity was used as a proxy for phage integration due to the rapid degradation of such elements in Rickettsiales genomes^21^) (Fig. 3e-f; Fig. S12). Although the decrease in anti-phage defense systems diversity seems to favor phage genomic integration, a late lytic cycle induction would still prove detrimental to the Rickettsiales host. An in-depth look into the Wolbachia wNo genome (circular chromosome, GCF_000376585) showed the existence of scattered syntenic blocks deriving from a prophage related to Wolbachia phage WO (KX522565) (Fig. 3h). These regions that most likely originated from one prophage were found to be flanked by transposase domains associated with insertion sequences (Fig. 3h). Thus, the high transposase abundances in intracellular lineages (where they are subjected to pervasive negative selection) (Fig. 3d; Fig. S13) may confer a selective advantage by engaging in prophage sequestration. This evolutionary strategy is consistent with Fan et al., 2019^22^ findings, which showed that insertion sequences may function as defensive mechanisms by suppressing dangerous invading genes. Such a transposase-mediated phage sequestration would allow intracellular specialists to increase gene flux between isolated populations, as acclimatization to the eukaryotic niche (e.g., survival in cytoplasmic membrane-bound vacuoles in Anaplasmataceae) likely decreases recombination frequency^23^ and hampers adaptation. Thus, Rickettsiales’ transition to an intracellular lifestyle was shaped by accelerated gene acquisition from their eukaryotic hosts and phage-mediated HGT events that facilitated the spread of the newly acquired genetic repertoire between allopatric populations. Mobile genetic elements and fast evolution rates^24^ allowed intracellular lineages to capture phages and erode them, recycling from their genetic cargo the parts that enhance niche adaptation. In some intracellular lineages (e.g., *Orientia tsutsugamushi*) extensive mobilome expansions caused genomic bloating, misleadingly reversing the characteristic genome shrinkage trend (Fig. 3c).

### Early diversification

Molecular phylogenetic work performed in the late 1970s^25^ revealed mitochondria’s bacterial ancestry and fueled the revival of a previous postulation that sought to explain eukaryogenesis through symbiosis^26^. The ‘90s brought fresh wind to the endosymbiotic theory through studies showing shared ancestry between Rickettsia and mitochondrial lines of descent^2,27^. This newfound sister-group relationship became orthodoxy and permeated mainstream evolutionary biology, where it endured mostly uncontested for two decades^28^.

Both Alphaproteobacteria and mitochondria radiations are notorious for harboring fast-evolving lineages^29,30^ that in phylogenetic reconstructions are represented as long-branched. As these types of taxa have the potential to attract each other regardless of true evolutionary history, a strategy focused on the evaluation and evasion of long-branch attraction artifacts was successfully employed (Figs. S14, S15). A Bayesian phylogeny (CAT-GTR) based on a concatenation of 24 proteins (6 416 aligned sites) resolved with high statistical support (pp=0.98) a sister-group relationship between Rickettsiales and mitochondrial genomes (Fig. 4a; Supplementary Text; Fig.S16). Although this phylogenetic relationship is not novel (as it was recovered in a prior study^29^) its validity has recently come under scrutiny^31,32^. In the competing evolutionary scenario mitochondria branch before the radiation of ‘known Alphaproteobacteria’^31,32^. The use of GC-rich mitochondria, the presence of a problematic taxon (MarineAlpha9 Bin5, GCA 002937595), and the existence of fast-evolving sites were all suggested as contributors to the Rickettsiales-mitochondria sister-group relationship^31,32^. The current study, however, favors this evolutionary model, as this phylogenetic relation remained stable after incremental removal of highly variable sites (present in the protein alignment), replacement of the disputed taxon, and usage of low GC mitochondrial genomes (median GC=34.67% ± 2%) (Figs. S16-S22). However, it is important to emphasize that in the ‘mitochondria outside known Alphaproteobacteria’ scenario Rickettsiales appear as the most basal and evolutionary closest bacterial lineage to the mitochondrial radiation (mitochondria forms a sister-group relationship with the lineage leading to Rickettsiales and thus to all ‘known Alphaproteobacteria’)^32^. The Rickettsiales-mitochondria sister-group relationship was further substantiated by the detection of mitochondrial-derived genes^33^ encoded in Rickettsiales genomes (Fig. 4b). These eight genes were found to be simultaneously present in very few genomes (37 out approx. 32 000) generally affiliated with ancient Rickettsiales and Rhodospirillales. This distributional restriction points towards vertical inheritance rather than HGT phenomena, as Rhodospirillales and Rickettsiales are phylogenetically related (both parts of the Alpha II supergroup; Supplementary Text; Fig. S15). An additional electron transport chain (ETC) phylogeny reiterated Rickettsiales/mitochondria sister-group relationship and indicated the presence of aerobic respiration capacity in the mitochondrial ancestor (Fig. S23). Though the sister-group phylogenetic connection is not surprising, its retrogression to ancient stage Rickettsiales provides a unique opportunity for inferring the metabolic landscape of the mitochondrial ancestor.

**Fig. 4.**
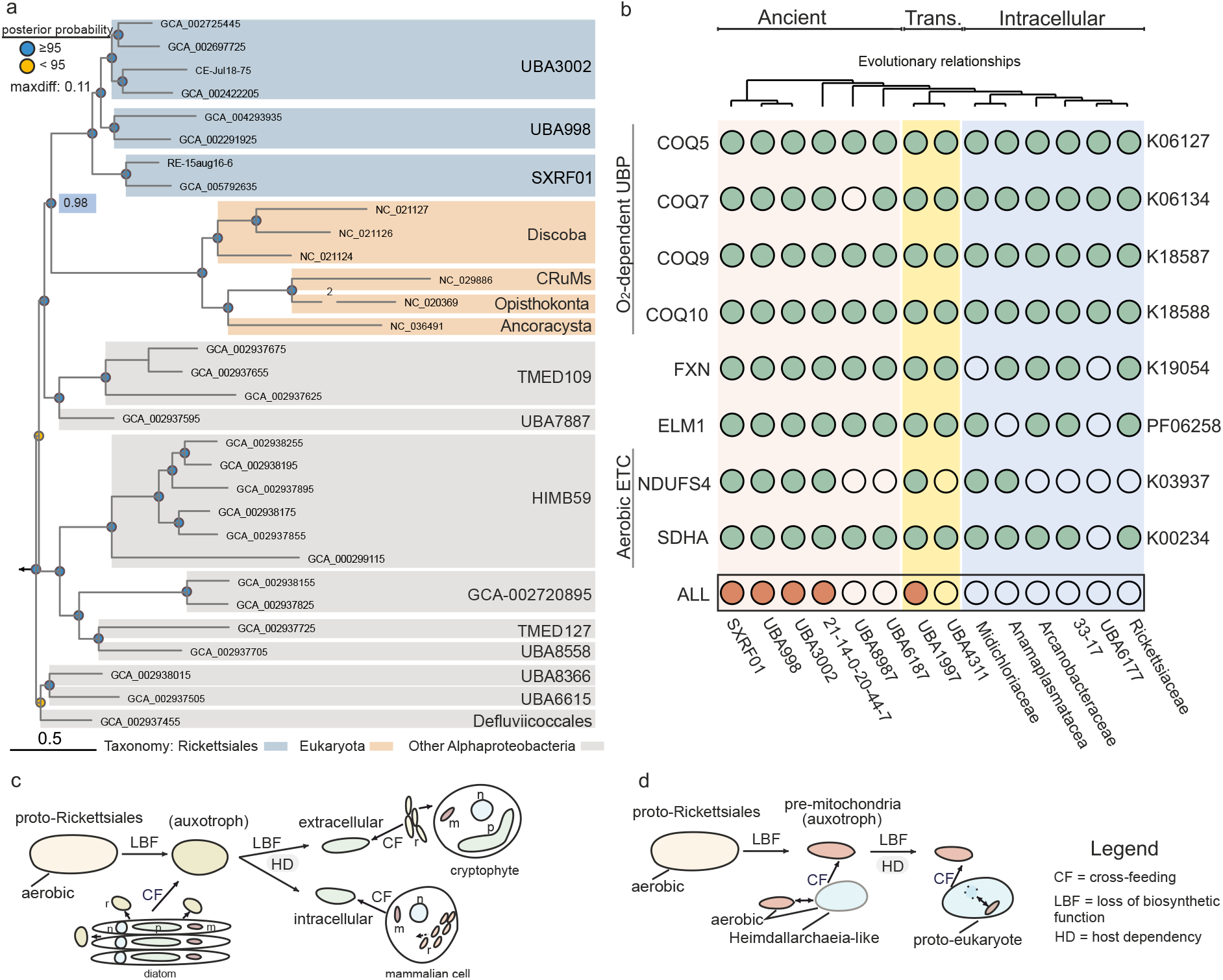
Rickettsiales-mitochondria evogenomic investigations. a) Mitochondrial-centric genome-based phylogeny generated through Bayesian inference (-cat -gtr -dgam4). Posterior probability values are represented through colored circles (top left legend). The posterior probability value (0.98) for the Rickettsiales/mitochondria node is highlighted with a blue box. The length of the fast-evolving Nuclearia simplex mitochondrial branch (NC_020369) is shown reduced (2× the scale bar). The scale bar indicates the number of substitutions per site. b) Mitochondrial-derived proteins/protein domains found in Rickettsiales families. Trans.: transition stage. O_2_-dependent UBP: Oxygen-dependent ubiquinone biosynthetic pathway; Aerobic ETC: Aerobic electron transport chain. COQ5: 2-methoxy-6-polyprenyl-1,4-benzoquinol methylase; COQ7: 3-demethoxyubiquinol 3-hydroxylase; COQ9: ubiquinone biosynthesis protein COQ9; COQ10: coenzyme Q-binding protein COQ10; FXN: frataxin; ELM1: mitochondrial fission ELM1; NDUFS4: NADH dehydrogenase (ubiquinone) Fe-S protein 4; SDHA: succinate dehydrogenase (ubiquinone) flavoprotein subunit. ALL: simultaneous presence of all eight genes in at least one genome. c-d) Schematic representation for cross-feeding-based scenarios for the evolution of Rickettsiales (c) and mitochondria (d).

Ancestral metabolism inference was based on metabolic pathways shared between the basal lineages of ancient stage Rickettsiales (SXRF01, UBA998, and UBA3002), as they encompass the order’s earliest evolutionary radiation to date. Against this backdrop, proto-Rickettsiales emerge as an oxygen-respiring free-living flagellated heterotroph that could detect (sigma factors and signal transduction domains) and react (aerotaxis, chemotaxis) to chemical signals encountered in a heterogeneous aquatic niche. This ancestral species gave rise to two radiations that took divergent evolutionary paths: while one brews dangerous pathogens (i.e. Rickettsiales lineage), the other gave birth to eukaryotes (i.e. mitochondrial lineage) (Fig. 4c-d). In the Rickettsiales lineage, the free-living bacteria started to exploit a new resource-rich niche comprising extracellular metabolites released by microbial eukaryotes (e.g. UBA3002 family). This newfound nutrient source favored the emergence of auxotrophic bacterial mutants^34^ that lost the ability to autonomously produce the already available biomolecules. As cross-feeding interactions strengthen, multiple metabolic pathways became redundant and susceptible to deleterious mutations (see the evolution of amino acid biosynthetic pathways Fig. 1b). The genomic shrinkage manifested in biosynthetic capacity reduction (e.g. loss of glyoxylate cycle and ED pathway), and pushed Rickettsiales towards obligate symbiosis (i.e. UBA1997). To avoid the eukaryotic partner loss and thus the vital supply of metabolites, Rickettsiales favored internalization (e.g. Anaplasmataceae). Inside nutrient-rich eukaryotic cells, bacterial metabolic degradation was accelerated through neutral ratchet-like gene losses^2^, thus giving rise to genomic architectures that hallmarked reductive genome evolution.

By drawing parallels between present-day genomic architectures of eukaryotic host-restricted bacterial symbionts^35,36^ and mitochondria^37^, it becomes evident that long-term mutualistic interactions favor function-centric genome degradation. Namely, the genomes of *N. deltocephalinicola* (0.11 Mbp) and *A. ciliaticola* (0.29 Mbp) bacteria were degraded towards the minimal metabolic potential needed to maintain the mutualistic symbiosis: biosynthesis of essential amino acids and anaerobic respiration, respectively^35,36^. As for mitochondria, their gene-impoverished genomes largely retained genes encoding for an aerobic electron transport chain employed by eukaryotes in respiration (albeit not exclusively). Thus, the bacterial metabolic circuitry that endures over long-lasting mutualistic interactions is likely reduced to keystone functions that are essential in sustaining the symbiotic partnership. By setting the phylogenetic relationship between Rickettsiales and mitochondrial ETCs into a ‘genome reduction to vital function’ framework, it becomes apparent that the vertically inherited aerobic respiration played a central function in the symbiotic interaction that fueled eukaryogenesis (as it was preserved throughout all evolutionary stages that led to modern-day eukaryotes). However, the emergence of aerobic respiration as a cornerstone in eukaryogenesis breaks away from the latest revisions of the syntrophy model^38^. In this scenario, formulated on metabolic pathways inferred to be present in the ancestor of Asgard superphylum (i.e. symbiotic partner of future mitochondrial lineage), eukaryogenesis emerged as a series of hydrogen-based syntrophic interactions set in a largely anoxic milieu. While the anaerobic nature of Asgard’s ancestor is conclusive, its role in eukaryogenesis is debatable as evolutionary reconstructions show eukaryotes branching from within an already differentiated Asgard lineage with facultative aerobic metabolism (i.e. Heimdallarchaeia)^39,40^. Though oxic eukaryogenesis departs from the hydrogen-based interactions of the syntropy model, it falls in line with the previously proposed ‘aerobic protoeukaryotes’^40^. In an extension of the latter, eukaryogenesis debuted with cross-feeding interactions between two microbial lineages undergoing an accelerated process of adaptation to a rapidly extending oxic environment. Here, the proto-Heimdallarchaeia (with mixed membrane lipids^41^) expanded its niche space by acquiring bacterial oxygen-dependent metabolic pathways^40^. This habitat expansion most likely pushed the archaeal lineage towards establishing new syntrophy to supplement the ones lost (from the previously anoxic niche). While it is impossible to pinpoint the nature of such cross-feeding interactions, speculative inferences regarding the type of possible trophic dependencies can be drawn from metabolic reconstructions of the closest phylogenetic clades (i.e. Heimdallarchaeia and SXRF01 Rickettsiales). In this context, proto-Heimdallarchaeia’s transition to an oxygenated environment was associated with the loss of its anaerobic B_12_ biosynthetic pathway. This inference stems from the persistence of B_12_-dependent enzymes in Heimdallarchaeia (e.g. methionine synthases and methylmalonyl-CoA epimerases/mutases), and the pathway’s presence in its anaerobic sister lineage^42^. To alleviate B_12_ auxotrophy, the proto-Heimdallarchaeia likely established cross-feeding interactions with producers extant in the oxygen-containing niche. One such candidate could have been proto-SXRF01 Rickettsiales (SXRF01 is the basal lineage of the ancient stage), a family comprised of lineages recovered from aquatic habitats and shallow lake sediments (i.e. the MAG ss-metabat2.31) with the capacity for de novo aerobic B_12_ synthesis. In such a cross-feeding partnership bacteria could provide the vital B_12_, while the archaea could deliver organics used to fuel the bacterial TCA cycle (and thus the aerobic respiration).

## Conclusion

Harnessing cultivation-independent genomics for charting diversity (i.e., genetic and functional) on phylogenomic frameworks enabled the unearthing of eco-evolutionary processes that steered harmless aquatic bacteria to intracellular parasites/symbionts. Rickettsiales’ evolutionary history could be diced into three chronological stages marked by specific genomic architectures and metabolic wirings. Ancient lineages were typified by a genomic/metabolic blueprint, usually associated with a heterotrophic free-living lifestyle that steadily degraded in the transition and intracellular stages which bore the hallmarks of reductive genome evolution. Genomic shrinkage was found to precede the advent of intracellularity and to be driven by the genetic drift favored by snowballing cross-feeding dependencies. The transition stage was marked by fast evolutionary changes (depicted by long branches in Fig. 1a) and lifestyle alterations. Here, extracellular lineages (i.e. UBA1997) prone to establish cross-feeding interactions with their eukaryotic kin found themselves steered by reductive genome evolution towards obligate intracellular specialization (i.e. UBA4311). The early radiations of the intracellular stage enhanced their niche adaptability by exploiting phages to propagate the newly acquired (and rewired) eukaryotic genetic material. Basal Rickettsiales were found to accommodate some of the, in very few bacterial lineages enriched, proteins expected to have been present in the mitochondrial ancestor. Evolutionary history reconstructions placed them as the closest relatives to modern mitochondria, thus providing the possibility to refine the metabolic landscape that favored the eukaryote’s emergence. In light of present findings, the dawn of eukaryogenesis debuted in a (micro)oxic aquatic habitat with two prokaryotic lineages likely engaging in cross-feeding interactions. Microbial genomes bear the marks of their evolutionary past. By viewing current bacterial diversity through an evolutionary genomics lens, it will become possible to grasp not just the eco-evolutionary drivers that shaped our distant past, but also those that may affect our future. Understanding how lifestyle transitions shape bacterial genomic architectures will likely open avenues for improved detection and surveillance of possible emerging diseases.

## Methods

### Assembly and binning

Raw Illumina metagenomic reads were pre-processed to remove low-quality bases/reads and adaptor sequences using the BBMap^43^ v36.1 package. Briefly, PE reads were interleaved by reformat.sh^44^ and quality trimmed by bbduk.sh^45^ (qtri =rl trimq=18). Subsequently, bbduk.sh was used for adapter trimming and identification/removal of possible PhiX and p-Fosil2 contamination (k =21 ref=vectorfile ordered cardinality). Additional inspections (i.e. *de novo* adapter identification with bbmerge.sh^46^) were performed to ensure that the datasets meet the quality threshold necessary for assembly. The pre-processed reads were assembled independently with MEGAHIT v1.1.5^47^ using the k-mer sizes: 39 49,69,89,109,129,149, and default settings. The pre-processed metagenomic datasets were mapped using bbwrap.sh^48^ (kfilter=31 subfilter=15 maxindel=80) against the assembled contigs (longer than 3 Kbp) in a sample-dependent fashion. The resulting BAM files were used to generate contig abundance profiles with jgi_summarize_bam_contig_depths (--percentIdentity 97)^49^. The contigs and their abundance files were used subsequently for hybrid binning with MetaBAT2^49^ (based on tetranucleotide frequencies and coverage data; default settings).

Post-binning curation was achieved by applying a taxonomy-based approach coupled with a GC cut-off. Briefly, the predicted proteomes (PRODIGAL^50^ v2.6.3) of individual bins were queried (using mmseqs^51^ search) against the curated prokaryotic GTDB database (R05-RS95)^52^. The obtained results were further converted into a BLAST-tab formatted file (using mmseqs convertalis) from which individual top hits (cut-offs: E-value 1e-3, identity 10%, coverage 10%, bitscore 50) were extracted, and their taxonomic labels used to decorate the queried proteomes. Taxonomic information was used to classify each bin at class level (i.e. Alphaproteobacteria) and to discard individual constituent contigs for which taxonomic homogeneity was not achieved (more than 30% of the taxonomic labels belonged to a different class). Contigs without taxonomy information or for which the GC content deviated by more than 15% from the bin median value, were discarded as well. Bin completeness, contamination, and strain heterogeneity were estimated by CheckM^53^ v1.1.3 (using the lineage_wf workflow). Bins with estimated completeness above 40% and contamination below 5% were denominated as metagenome-assembled genomes (MAGs). All the obtained MAGs were taxonomically classified with GTDB-Tk v1.4.0 (database release R05-RS95) using default settings^52^. The MAGs belonging to Rickettsiales were selected to be used in the present study (n=49). The obtained dataset was supplemented with 80 publicly available genomes (contamination below 5%).

### Rickettsiales phylogenomics

To obtain a broader view of Rickettsiales genomic landscape we recovered all representative ‘species’ present in the curated GTDB R05-RS95 release (comprising 191 527 bacterial genomes organized into 31 910 ‘species’ clusters)^52^. For methodological details regarding species-level clustering and representatives’ selection see Parks et al., 2020^54^. The obtained genomes were screened for the presence of collocated, lineage-specific marker sets by CheckM^53^ v1.1.3 (using the lineage_wf workflow) to establish their redundancy/contamination. Rickettsiales genomes with less than 5% redundancy/contamination (n=80) were used together with the obtained MAGs (n=49) for downstream analyses.

Phylogenies were crafted around a set of 57 proteins that were previously evaluated as being phylogenetically informative and suitable for deep evolutionary inferences^55,56^. These proteins are commonly subjected to different degrees of negative selection (hence the sequence conservation), and thus their amino acids exhibit site-specific variations in their substitution rates. Phylogenies constructed under site-heterogeneous replacement models were used to account for this between-site variation in the substitution process. Outgroup rooting was performed to confer directionality to phylogenomic trees. For outgroup determination, an evolutionary reconstruction was made between selected Rickettsiales (n=3) and Alphaproteobacteria members (n=32) (Fig. S3). Genomes belonging to slow-evolving Alphaproteobacteria clades were selected to mitigate the potential long-branch attraction (LBA) caused by fast-evolving lineages and Rickettsiales (as long branches cannot attract each other when there is only one such branch in the analysis). PRODIGAL^50^ v2.6.3 (default settings) was used to predict protein-coding genes for each genome/MAG (n=35). The obtained proteomes were scanned with HMMER hmmscan^57^ v3.1b2 (with an E-value threshold set at 1E-10; -prcov 70 -hmcov 70) against a locally installed TIGRFAMs^58^ (v15.0) database. 57 ubiquitous single-copy protein sequences (see Supplementary Data S1) were extracted from the genomes (based on TIGRFAMs accession numbers). Phylogeny-aware multiple sequence alignments (MSAs) were constructed for each phylogenomic marker (n=57) using the software PASTA^59^ v1.12 with default settings. The obtained alignments were trimmed with BMGE^60^ v1.12, using the -g 0.5 setting, to retain aligned regions suitable for phylogenetic inferences. The different MSAs were concatenated in a supermatrix (21 047 aligned sites) prior to performing genome-focused phylogenies. As Alphaproteobacteria phylogenies are plagued by LBA phenomena^29^ they were generated through Bayesian inference with site-specific profiles, as this framework was found to be robust against such artifacts^61,62^. Thus, four independent chains were run through PhyloBayes MPI^63^ 1.8b with: i) site-specific equilibrium frequency profiles (-cat option), ii) uniform exchangeabilities (-f81 option), and iii) rate variation across sites modeled through a discretized gamma distribution with 4 categories (-dgam 4). The chains were stopped after 22 252 trees (for each chain) and their burn-in (i.e. number of points before the chain has reached stationarity) was estimated at 2 300 points by Tracer^64^ v1.7.1. The convergence of parameters and tree space between the 4 chains was assessed by tracecomp and bpcomp software (as implemented in PhyloBayes MPI^63^ 1.8b). The obtained values for effective sizes (> 1 400), discrepancies (< 0.16), and the maximum difference in bipartition frequencies (= 0.11) indicated convergence. Thus, a consensus tree was calculated by pooling the trees (n=79 800) from the 4 chains.

The curated Rickettsiales genomic dataset (n=129) was subjected to taxonomy-centered downsampling (as the size of the dataset was larger than recommended optimum) (PhyloBayes MPI manual^63^). Thus, 53 genomes spanning 14 family-level clades were selected for evolutionary reconstructions. Phylogenetic marker identification (n=57) and alignment was performed as previously described. The obtained supermatrix (18 380 aligned sites) was used to generate 3 phylogenies under site-heterogeneous models, as the exclusion of fast-evolving lineages was unavoidable. The following models were used under PhyloBayes MPI^63^ 1.8b:

1. i) empirical profile mixtures (-catfix C60), ii) data-inferred exchangeabilities (-gtr option), and iii) rate variation across sites modeled through a discretized gamma distribution with 4 categories (-dgam 4). 4 independent chains.
2. i) site-specific equilibrium frequency profiles (-cat option), ii) uniform exchangeabilities (-f81 option), and iii) rate variation across sites modeled through a discretized gamma distribution with 4 categories (-dgam 4). 4 independent chains.
3. i) site-specific equilibrium frequency profiles (-cat option), ii) data-inferred exchangeabilities (-gtr option), and iii) rate variation across sites modeled through a discretized gamma distribution with 4 categories (-dgam 4). 4 independent chains.

The obtained phylogenies were evaluated for: i) convergence in tree space (maximum difference in bipartition frequencies < 0.3 for ≥ 3 chains), ii) discrepancies among chains (< 0.3 between ≥ 3 chains) and iii) effective sample size (> 100 between ≥ 3 chains). These assessments were performed using the tracecomp and bpcomp software. The most accurate phylogenomic reconstruction (between the ones achieving the up-mentioned criteria) was designated based on goodness-of-fit tests on site-specific amino acid usage patterns (using readpb_mpi as implemented in PhyloBayes MPI^63^ 1.8b). Consequently, a consensus CAT-F81 phylogeny (192 240 trees; 3 chains; maxdiff=0.2) was selected to depict the evolutionary history of Rickettsiales.

### Genome annotations

Protein-coding genes were predicted with PRODIGAL^50^ v2.6.3. Protein domains were annotated by querying the obtained proteomes (using the hmmscan-based ‘pfam_scan.pl’ script) against the HMM database present in Pfam^65^ release 32. Additional domain architectures and protein annotations were performed by running InterProScan^66^ (v5.24-63.0) with the databases CDD (v3.14), SMART (v7.1), and HAMAP (v201701.18), respectively. Protein annotation space was further enlarged by running hmmsearch (-evalue 1E-7 -prcov 70-hmcov 70) against COGs^67^ and TIGRFAM^58^ HMM databases. BlastKOALA^68^ was used to assign KO identifiers to orthologous genes. The obtained K numbers were further mapped to KEGG pathways and modules by the MetQy^69^ R package. Transmembrane topology and signal peptide prediction were performed by Phobius^70^ v1.01 with default settings. Structures were predicted for selected proteins using the Phyre2 web portal^71^. The locations of ribosomal RNA genes were predicted using barrnap 0.9. Rigorous manual curation was applied to ensure metabolic pathway completeness. The GTDB R05-RS95 database (approx. 32 000 genomes) was annotated with protein domains and KO identifiers using the Pfam^65^ release 32 and KOfam^72^, respectively. Carbohydrate-active enzymes were identified in 129 Rickettsiales genomes/MAGs by run_dbcan v2.0.11 software^73^. CRISPR-Cas systems were identified and classified by CRISPRcasIdentifier^74^ v1.1.0 with default settings. Innate anti-phage systems and other defense mechanisms were recognized by using custom searches against the annotated genomes/MAGs. Additional anti-phage defense mechanisms were predicted by using DefenseFinder^75^ v1.2.1 software with default settings. The detection of biosynthetic pathways of secondary metabolite production was performed by antiSMASH^76^ v4.2.0 software (default settings). Identification of prophages among the genomes/MAGS was achieved by PhiSpy^77^ v4.2.19 software (--phage_genes 3 option). Ankyrin gene-context analysis was performed on the genomes/MAGs containing contigs with a minimum of 5 genes on each side of a protein having at least one Ank repeat (Ank, Ank_2, Ank_3, Ank_4, Ank_5) (n=264). The probability of finding any domain by chance in a random subset of ten genes was calculated (for each genome) using the hypergeometric distribution^78^ (without replacement) in R with the function phyper (stats package). P-values were corrected using the Benjamini-Hochberg procedure^79^ to account for type I errors arising from multiple comparisons (q-values < 0.01).

Rickettsiales families with intracellular lifestyles (i.e. Midichloriaceae, Anaplasmataceae, Arcanobacteriaceae, and Rickettsiaceae) were found to form a monophyletic unit that distinguished itself through an increased number of transposase-coding genes. 1 960 proteins annotated as transposases (from these four family-level clades) were used in orthogroup inference by OrthoFinder^80^ v.2.5.2 (-I 3 -S blast). Orthogroups composed of a minimum of five sequences (n=44) were used for codon alignments with prank^81^ v.170427 (-codon -F). Site-specific selection pressure was inferred based on synonymous and nonsynonymous substitution rates as implemented in FUBAR^82^ (HYPHY 2.5.32) software. ISEScan (https://github.com/xiezhq/ISEScan) was used to scan for putative insertion sequence elements, both with -removeShortIS option and without to get all elements and only full-length ones respectively. The target GCF_000376585 genome was further scanned against the latest version of the Isfinder^83^ database using local blastn^84^ with default parameters.

### Catalyzed reporter deposition-fluorescence in situ hybridization (CARD-FISH) and image analysis

Two oligonucleotide probes were designed to target the 16S rRNA sequences of the UBA1997 and UBA3002 families. 16S rRNA genes extracted from MAGs were aligned using SINA web aligner^85^ and added to the SILVA^86^ 138 SSU Ref NR 99 database using the ARB^87^ software. After manual alignment curation, an x100 bootstrapped tree was constructed with the Maximum Parsimony Randomized Accelerated Maximum Likelihood algorithm (RaxML) and GTR-GAMMA model^88^. For both Rickettsiales families, the probe candidates were retrieved and tested with the Probe_Design and Probe_Match functions in ARB^87^. The promising oligonucleotides were additionally checked using the online TestProbe tool^86^ with and without mismatches. Formamide concentrations for CARD-FISH hybridization buffers were estimated with the MathFish^89^ software. Designed probes labeled with horseradish peroxidase (HRP) were used to access the spatio-temporal dynamics of the targeted Rickettsiales families in a one-year data set collected from Řimov reservoir (CZ)^90^. In brief, water samples were collected at 5 stations along the longitudinal transect of the Řimov reservoir at three weeks intervals during the year 2015 (n=225). At stations with depths above 10 m, additional samples were taken from deeper strata. Water samples were fixed with formaldehyde on the site (2% final concentration) and transported to the laboratory within two hours. The filters for CARD-FISH analysis were prepared on the same day from 4 to 12 ml of water following the bacterial densities. For this purpose, water was filtered onto 47 mm diameter 0.2 μm pore-sized polycarbonate white membranes (Millipore, Merck, Darmstadt, DE), which were stored at −20°C for future analysis. Subsequently, filters were embedded in 0.1% agarose and enzymatically treated with lysozyme (10 mg ml-1, 60 min, 37°C) and achromopeptidase (60 U ml-1, 30 min, 37°C) to permeabilize the bacterial cells. Hydrogen chloride (0.01 M, 10 min) was used to deactivate naturally present peroxidases. After this pretreatment, the filter sections were hybridized for 2 h at 35°C. The concentration gradients within the 20% range based on MathFISH results were used to select the optimal formamide concentration for the hybridization buffer. After the standard washing procedure, fluorescein-labeled tyramines were used for signal amplification (30 mins, 37°C). All processed samples were counterstained with 4′,6-diamidino-2-phenylindole (DAPI) and analyzed by epifluorescence microscopy (Zeiss Imager Z2, Carl Zeiss, Oberkochen, DE) with a colibri LED system and the following filter sets: DAPI 49 (Excitation 365; Beamsplitter TFT 395; Emission BP 445/50), fluorescein 38 HE (Excitation BP 470/40; Beamsplitter TFT 495; Emission BP 525/50) and chlorophyll A 62 HE (Excitation BP 370/40, 474/28, 585/35; Beamsplitter TFT 395 + 495 + 610; Emission TBP 425 + 527 + LP615 HE). The proportions of hybridized cells were calculated manually for at least 500 DAPI-stained cells. The image sets including DAPI, autofluorescence, and fluorescein channel images were prepared for the estimation of the width and length of the hybridized cells. The dimensions of 100 individual probe-positive cells were measured on the DAPI images with the help of NIS – Elements AR v4.6 software using the Annotation and Measurements tool.

UBA1997-positive filters were further used for double hybridization as previously described^91^. In these cases, the Alexa568 labeled tyramides were used for amplification of hybridization signal from UBA1997 cells, and fluorescein-labeled tyramides were applied after hybridization with Crypto B probe (targeting 74.3% of available Cryptophyceae 18S rRNA sequences^92^). Alexa568 signal was accessed with the 62 HE filter set described above. Three-channel z-stacks were recorded with an Axiocam 506 (Carl Zeiss, Oberkochen, DE) using a 63x plan-apochromat objective. The merging of images was performed in the ZEN v2.6 software (Carl Zeiss, Oberkochen, DE) using the focus extension function.

### Mitochondrial phylogenomics

The most recent (and generally accepted) views on the identity of the pre-mitochondrial lineage are divided into two scenarios. One in which the mitochondrial ancestor branches outside Alphaproteobacteria, and another one in which it branches within (near Rickettsiales). The phylogenetic position of mitochondria was assessed by applying the “long-branch extraction method”, inspired by Siddall and Whiting^93^ to phylogenies estimated by Bayesian inference (with site-specific equilibrium frequency profiles, CAT-GTR). An Alphaproteobacteria phylogeny was generated using the slow-evolving lineages described in Fan et al.,^29^. This phylogeny was used as a backbone as it is not affected by LBA phenomena (as there are no long branches to attract each other). The placement of Rickettsiales was evaluated by generating a phylogeny between this fast-evolving lineage and slow-evolving Alphaproteobacteria ones since long branches cannot attract each other in their mutual absence. The same rationale was followed for the phylogenetic placement of mitochondria. To estimate if the fast-evolving Rickettsiales and mitochondria attract each other they were used together (with the backbone taxa) in a phylogenomic reconstruction. All obtained topologies were evaluated (Fig. S14).

The generated phylogenies (n=4) showed stability in the topological placement of the slow-evolving Alphaproteobacteria lineages, as the presence of fast-evolving ones did not affect the backbone structure of the trees. Rickettsiales and mitochondria were both placed independently in the slow-evolving Alpha II group (Fig. S14). When present together, Rickettsiales and mitochondria showed a sister group relationship that branched within the same Alpha II group. The two fast-evolving lineages displayed the same topological placement in the absence of the other as together, rendering their phylogenetic proximity an evolutionary linkage rather than an LBA phenomenon.

Precise evolutionary placement of the mitochondrial lineage was accomplished by evaluating the phylogenetic relationships between bacterial taxa related to the Alpha II group. To this end, a set of 19 bacterial genomes used by Fan et al.^29^ was supplemented with 8 basal Rickettsiales and 6 protist mitochondrial ones (Supplementary Data S1). Protein coding genes were predicted by PRODIGAL^50^ v2.6.3 software (default settings). The MitoCOGs protein dataset was downloaded from https://ftp.ncbi.nih.gov/pub/koonin/MitoCOGs/. From this, sequences affiliated with 24 MitoCOGs were extracted and clustered with mmseqs^51^ (easy-cluster -min-seq-id 0.85 -c 0.8 -cov-mode 0). Each protein-containing cluster was independently aligned with MAFFT-L-INS-I^94^ v7.471. Hidden Markov models (HMM) were constructed using HMMER v3.3 (hmmbuild) from the obtained alignments. HMM-specific e-values were determined by assessing the e-values for the target (as present MitoCOGs) and non-target hits. The mitochondrial (GC=34.67% ± 2%) and bacterial genomes/MAGS (n=33) were screened using hmmsearch (--domblout) for the presence of the 24 MitoCOG markers. The identified proteins were independently aligned with PASTA^59^ v1.12 (--estimator=raxml) and trimmed with BMGE^60^ v1.12 (default settings). The trimmed alignments were concatenated (6 416 aligned sites) and used in phylogenetic reconstructions. Two phylogenies were generated (PhyloBayes MPI^63^ 1.8b; 4 chains/analyses) under CAT-F81 and -GTR models. The obtained phylogenetic reconstructions were evaluated as previously described. As both phylogenies displayed similar topologies, the CAT-GTR one was shown (18 064 trees; 4 chains; maxdiff= 0.11) as it modeled better the site-specific amino acid propensities (z-score=2.86). Site-specific evolutionary rates were calculated using IQ-TREE^95^ v 2.1.3 with the following settings: -m LG+F+R10 -n 0 -wsr. The topmost rapidly evolving sites were incrementally removed from the supermatrix with SiteStripper^96^ v1.03 (-rf iqtree -f 0.95) to alleviate the possible systematic errors and/or substitution saturation.

### Statistics

All statistics were performed with R^97^ v4.0.3 and RStudio^98^ v1.3.1093, Apricot Nasturtium software. Data normality was assessed by the Shapiro-Wilk test, followed by residues distribution visualization. Simple comparisons between two datasets were performed by student t-test whenever data was following normal distribution and Wilcoxon rank sum test for non-normal datasets. Parametric analysis of variance (ANOVA) was performed for normally distributed datasets followed by multiple pairwise comparisons with Tukey’s test. Non-parametric datasets analysis of variance was performed by using the Kruskal-Wallis test followed by pairwise comparisons with pairwise Wilcoxon rank sum tests. For non-normal samples, the correlation index was assessed using the Pearson-rank correlation test. Variable interaction as well as multiple correlations were assessed by multiple regression models. Discussed tests were performed using the corresponding functions within the stats v.4.1.3 package (part of R).

## Acknowledgments

We are grateful to Petr Znachor, Pavel Rychteck, Petr Porcal, Thomas Posch, Eugen Loher, the Canton of Bern’s Laboratory for Water and Soil Protection (GBL), and the crew of the research vessel “Kormoran” (LUBW-ISF Langenargen) for their support during sample collection. We also want to thank Alizée Le Moigne for her assistance with the statistical analyses.

## Author contributions

Conceptualization: A.-S.A. with involvement from L.S.M. Sampling, sample processing: A.-S.A., T.S., V.S.K., S.-J.P. Bioinformatics: A.-S.A., L.S.M., T.S., V.S.K., P.-A.B., M.-C.C., S.-J.P., R.G. Statistics: L.S.M. CARD-FISH: T.S., I.M. Writing, original draft: A.-S.A. with input from L.S.M. Writing, review, and editing: L.S.M, T.S., V.S.K., P.-A.B., M.-C.C., S.-J.P., I.M., R.G. Funding acquisition: A.-S.A.

## Data and materials availability

All sequence data generated during this study have been deposited in the EBI/NCBI (Bioprojects: PRJEB35770, PRJNA429145, PRJEB56877, PRJEB56878, and PRJNA279271). The generated data supporting the conclusions of this study may be found at figshare: 10.6084/m9.figshare.21269577. All additional important data supporting the study’s conclusions are included in the publication and its supplemental material files.

