## Supplementary Materials for "Rickettsiales’ deep evolutionary history sheds light on the emergence of intracellular lifestyles"

**The PDF file includes:**

Supplementary Text
Figs. S1 to S28
Tables S1 to S4
References

**Other Supplementary Materials for this manuscript include the following:**

Data S1

**Supplementary Text**

<u>Sampling sites</u>

Samples from five freshwater lakes (that range in trophic status from oligotrophic to eutrophic; the Czech Republic and Switzerland), a cave pool (Switzerland), and shallow lake sediments (Republic of Korea) were used to recover genomic information from Rickettsiales present in diverse freshwater niches. The obtained genomic dataset was augmented with a metagenomic profile of the Caspian Sea (brackish environment), and publicly available data generated from various environments (as recovered from GTDB R05-RS95 that spans 194 600 genomes organized into 31 910 species clusters).

Římov Reservoir (470 m a.s.l., 48°50'N, 14°29'E, Czech Republic) is a meso-eutrophic, canyon-shaped dimictic water body with an area of 2.0 $km^2$ (length 13.5 km, the volume of 34.5 × 106 $m^3$, mean water retention time 77 days, maximum depth of 43 m) that was built during 1974–1979 by damming a 13.5 km long section of the River Malše. The sampling was performed between June 2015 and August 2017, above the deepest point of the reservoir by using a Friedinger sampler. 20 L of water were collected from 0.5 (n=10) and 30 m (n=8) depths and subjected to sequential peristaltic filtration through a series of 20, 5, and 0.2-µm-pore-size polycarbonate membrane filters (⌀ 142 mm) (Sterlitech Corporation, USA). The sample collection and filtration steps were similar for the rest of the lakes/pools unless otherwise stated. Jiřická pond (892 m a.s.l., 48°36.96'N 14°40.59'E, Czech Republic) is a dystrophic humic water body with an area of 0.035 $km^2$ (volume 6.59 x103 $m^3$, mean water retention time 9 days, maximum depth of 3.7 m), located in the Novohradské mountains of Southern Bohemia. Five epilimnia (0.5 m depth) water samples were collected between May 2016 and August 2017. Lake Zurich (406 m a.s.l., 47°18'N, 8°34'E, Switzerland) is an oligomesotrophic, perialpine monomictic water body, with an area of 67.3 $km^2$ (length 40 km, volume 3.3 $km^3$, mean water retention time 1.4 years, maximum depth of 136 m). Nine water samples were collected between 2013 – 2018 from the epilimnion (5 m depth, n=5) and hypolimnion (80/120 m depth, n=4) layers, and processed as described above. Lake Thun (558 m a.s.l., 46°41'N, 7°43'E, Switzerland) is an oligotrophic, alpine water body with an area of 48.3 $km^2$ (length 17.5 km, volume 6.5 $km^3$, mean water retention time 1.8 years, maximum depth of 217 m). Two water samples were collected in June 2018 from 5 and 180 m depths. Lake Constance (395 m a.s.l., 47°32'N, 9°31'E, Swiss Confederation) is an oligotrophic perialpine lake with an area of 473 $km^2$ (length 63 km, volume 48 $km^3$, mean water retention time 5 years, maximum depth of 252m). Four samples were collected in July and October 2018 from 5 m and 200 m depths.  Susan Reservoir (50 m a.s.l., 33°28.22'N, 126°23.22'E, Jeju, Republic of Korea) is an oligotrophic water body constructed in 1960 that is used for agricultural irrigation. One shallow sediment sample was collected in March 2008 at a depth of 0.5 m using a Petite Ponar dredge (Wildco, Saginaw, MI, USA). NGIII (591 m a.s.l., Switzerland) is a rock pool in the frequently flooded part of Bärenschact cave, situated 850 m below the entrance. The pool (average depth 0.4 m, surface approx. 9 $m^2$, volume approx. 3.5 $m^3$) is located at approximately half a kilometer distance from North Sump in the branch of the 'GaIery du Nord'. It is flooded when the karst water table of the sump rises higher than 28.5 m above its regular level. One water sample was collected in November 2018. More information about the Bärenschact cave can be found in Shabarova et al., 2014[1]. Entrance coordinates can be provided upon request (due to safety reasons).

DNA was extracted from the 0.22-µm filters (0.2- to 5-µm fraction) and sediments using the ZR Soil Microbe DNA MiniPrep kit (Zymo Research, Irvine, CA, USA) following the manufacturer's instructions. The total quantity of DNA was estimated using the Qubit dsDNA BR assay kit (Life Technologies, Foster City, CA, USA) on a Qubit 2.0 fluorometer (Life Technologies). DNA integrity was assessed by agarose gel (1%) electrophoresis and SYBR green I stain. Shotgun sequencing was performed using the Novaseq 6000 sequencing platform (2 × 150 bp) (Novogene, Hong Kong, China).

For the Caspian Sea, we made use of a previously available depth profile (15, 50, and 150 m) generated in October 2013, near Babolsar, Iran (52°36'E, 36°51'N). Details regarding sampling, physicochemical parameters, DNA extractions, and sequencing have been presented in the original study[2]. In this case (Caspian Sea samples), we reassembled and re-binned the publicly available sequencing data (SRR2026986, SRR2027816, and SRR2027830).

Rickettsiales environmental distribution as assessed by 16S rRNA gene fragments

The Rickettsiales genomes database (comprising 129 genomes and MAGs, see supplementary Data S1) was screened with barrnap version 0.9 (--kingdom bac) to detect rRNA genes present in each genome/MAG. 85 16S rRNA sequences (median length 1 495 bp) belonging to 12 Rickettsiales families were recovered and used to augment the SILVA[3] database release 138 (Ref NR 99). The 16S rRNA genes received the taxonomic strings from the genomes of provenance (previously classified by performing whole-genome phylogenies with GTDB-Tk[4] v1.4.0 software and R05-RS95 database). The SILVA database was altered through the replacement of present Rickettsiales sequences with the ones recovered from the genomic dataset. This was performed to i) link the physiological and ecological inferences derived from genome-centric analyses to environmental distribution data, ii) assess the contribution of environmental Rickettsiales to prokaryotic community structure, and iii) avoid some erroneous taxonomic assignments present in the database (see below).

To evaluate Rickettsiale's environmental distribution, 212 aquatic metagenomic datasets were screened (surface freshwater n=41, subsurface freshwater n=46, brackish n=13, marine n=96, and sediments n=16, supplementary Data S1). The analysis was centered on aquatic habitats since 95.8% of non-host-associated Rickettsiales genomes (n=23) were recovered from marine or freshwater environments (as shown by GTDB; see supplementary Data S1). Briefly, pre-processed Illumina datasets were converted to FASTA format and subsampled to 10 million sequences using reformat.sh[5]. These subsets were queried against the SILVA SSU database to identify RNA-like sequences by using MMSeqs2[6] with an E-value cutoff of 1E−3. The *bona fide* 16S rRNA gene sequences (as identified by SSU-ALIGN[7]) were further compared by blastn[8], in nucleotide space (E-value cutoff 1E−5), against the SILVA database amended with 16S rRNA genes recovered from Rickettsiales genomes. Taxonomic classification was performed for sequences that simultaneously had identity values ≥80% and alignment lengths ≥90 bp (sequences failing these thresholds were not used for downstream analyses). The taxonomic affiliation of each identified 16S rRNA read was inferred based on its best blastn hit. The relative abundances of Rickettsiales families were calculated as a percentage of the total 16S rRNA reads. The significance of family-level richness variation between habitats was estimated using non-parametric tests (i.e., Kruskal-Wallis and Dunn's).

The 16S rRNA sequences, used to augment the SILVA database, were uploaded into the SINA[9] Search and Classify service (https://www.arb-silva.de/aligner/; LCA classification based on taxonomies hosted by SILVA) to integrate the genomic-derived taxonomic labels into the larger 16S rRNA-based bacterial taxonomy. The attempt to link the used labels with

SILVA-derived ones was unsuccessful since the 16S rRNA sequences recovered from the genomic dataset (that did not belong to families with cultivated representatives) underperformed in taxonomic resolution. This outcome was a consequence of the reduced similarity between the recovered 16S rRNA sequences and the ones present in the SILVA database (Supplementary data). Surprisingly, the sequences belonging to the family UBA1997 were classified as mitochondrial (Supplementary Fig. S1). Since non-target filtering (i.e., removing sequences classified as mitochondria or chloroplast) is a common practice in prokaryotic environmental surveys[10–12], likely sequences belonging to UBA1997 are usually excluded from such studies.

212 metagenomic datasets recovered from aquatic habitats that span a broad range of trophic statuses (from oligotrophic to eutrophic), salinities (from freshwater to brackish and saline), and oxygen milieus (from oxic to microoxic) were used to assess the presence of Rickettsiales in environmental samples. Metagenomic-derived 16S rRNA reads were classified at the family level to circumvent the limited phylogenetic resolution associated with short sequence lengths. It must be pointed out that despite this inherent limitation, the taxonomic profiles obtained using 16S rRNA reads recovered from metagenomes routinely outperform the amplicon-based ones[13]. Phylogenetic signal belonging to Rickettsiales was detected in 88.8% of the analyzed samples, albeit at very low amplitude (median abundance 0.05%). This low abundance baseline was disrupted by high sporadic peaks in freshwater and brackish habitats (maximum abundance of approx. 3%). The family-level groups that registered high abundances (i.e., UBA1997 and UBA6187) were found not to relate to Rickettsiales cultured diversity (Fig. S2). Even though transient abundance peaks point towards the presence of a dynamic niche, the fact that family UBA1997 reached multiple times high densities (up to 2.7%) in distinct freshwater samples points towards habitat association. As higher diversity and abundances were recorded in brackish and freshwater habitats (Fig. S2), these low-salinity environments likely acted as diversification hubs for the present Rickettsiales radiation.

Rickettsiales phylogenomic trees

The Bayesian phylogenomic analyses reached convergence (maxdiff $< 0.3$ for $\geq 3$ chains) for the C60-GTR and CAT-F81 models. The CAT-GTR chains did not converge and were stopped after 100 000 trees per chain. However, when evaluated in a two-by-two fashion the chains showed low differences in bipartition frequencies (0.03 and 0.23, respectively). Upon scrutiny, the two consensus phylogenies (generated from two chains each) showed identity in the family-level branching patterns (Fig. S4). The discrepancy was found to lie in the internal branching of the UBA1997 family, and not in its phylogenetic positioning. Since the main objective of the analysis was to retrace the deep evolutionary history of the order (and to a lesser extent intrafamily relationships), the obtained phylogenies were further taken into consideration. Posterior predictive analyses on site-specific amino acid usage patterns showed that CAT-based models outperformed C60 in estimating the observed diversity (z-scores of 5.4/7.5 vs. 25.2 for C60). Under topological scrutiny, all 3 models returned very similar evolutionary histories (Fig. 1; Fig. S4). The only noteworthy difference was the position of the family Arcanobacteraceae, which appeared as a sister group of family 33-17 (in C60-GTR) or ancestral to UBA6177 and Rickessiaceae families (in CAT-GTR), albeit with low statistical support. This lineage was confidently placed with high statistical support in the CAT-F81 phylogeny (posterior probability of 1), which was further used to depict the evolutionary history of the order (Fig. 1).

Metabolism

Additional genome-inferred metabolic reconstructions revealed that most of the detected Rickettsiales families may be able to exploit the non-oxidative pentose phosphate pathway by transforming imported sugars (e.g., semiSWEET transporters) into glyceraldehyde 3-phosphate that could be further broken down to pyruvate (in the glycolysis core module; glyceraldehyde 3-phosphate to pyruvate). After decarboxylation, pyruvate is likely oxidized, losing one carbon while generating acetyl-CoA. This compound can further be used in the downstream biosynthetic pathways as well as the tricarboxylic acid cycle and/or the glyoxylate one in the SXRF01, UBA998, and UBA3002 families. Interestingly, ancient stage Rickettsiales had two-compound systems that may allow them to detect changes in both phosphate and nitrogen levels. The phosphate limitation two compound system (PhoPR) is composed of a phosphate regulon sensor histidine kinase (PhoR) and two regulators (PhoB-phosphate regulon response regulator and PhoP- alkaline phosphatase synthesis response regulator) that induce gene expression when phosphorus concentration falls. Additionally, ancient stage Rickettsiales families (SXRF01 and UBA3002) harbored the NtrB/NtrC two-compound system. NtrB is a nitrogen regulation sensor histidine kinase able to detect changes in intracellular 2-ketoglutarate to glutamine ratios (nitrogen availability indicator) and to further activate NtrC. Once activated NtrC can induce transcription of amtB ($NH_3$ transport) and glnALG ($NH_3$ assimilation) operons in SXRF01 or dppABC (dipeptide transport) in the UBA3002 family. The presence of both phosphorous and nitrogen two-compound systems in ancient stage Rickettsiales points towards capacity for gene expression regulation in response to fluctuating physiological or niche conditions. Another feature highlighting environmental plasticity among recovered Rickettsiales is the presence of multiple terminal oxidases exhibiting a wide range of oxygen affinities (from the low-affinity A1 to the high-affinity bd-I). Such oxidases can confer bacteria the capacity to adapt to niches of fluctuating oxygen concentrations. The existence of the cytochrome c oxidase cbb3-type (i.e., type C) in the ancient and transition stages likely facilitated the colonization of microoxic environments.

Considering their metabolic potentiality, the ancient stage Rickettsiales appear to be able to fulfill the needs of an independent heterotrophic lifestyle centered on the oxidation of amino acids and sugars. Their increased capability for proteinogenic amino acids biosynthesis together with the presence of sensorial/gene regulatory capacity (see Fig. 1; Tables S1-S2) and multiple terminal oxidates (with contrasting oxygen preferences) render them capable of surviving in dynamic niches. Whereas ancient and intracellular Rickettsiales showcase contrasting life strategies as inferred from their genomic properties and metabolic capacity, the transition stage appeared to bridge both. By sharing similar genomic properties with the intracellular stage (i.e., Fig. 2, GC% and estimated CDS) and exhibiting more ancient-like metabolic capabilities (i.e., proteinogenic amino acid biosynthesis capacity, cytochrome c oxidases cbb3 type), transition stage representatives appear to fill the gap in the evolution of Rickettsiales lifestyle strategies. Thus, by coupling sensible phylogenesis with genome-wide metabolic reconstructions it was possible to unveil stages and processes that led to emergences of parasitism in Rickettsiales bacteria.

Protein domains co-occurrence

The taxonomic milieu of selected protein domains co-occurring in the proximity of ANK-encoding genes was explored through screening of 31 910 prokaryotic genomes recovered from GTDB R05-RS95 database (in-house annotation) and online searches performed (June 2022) within the Pfam 35.0 database (https://pfam.xfam.org/)

1. IQ (PF00612) domain

- Distribution in Bacteria (GTDB R05-RS95): 64 sequences in 54 species.
- Distribution in Bacteria (Pfam 35.0 database): 21 sequences in 17 species.
- Distribution in Eukarya (Pfam 35.0 database): 39 318 sequences in 1 524 species.

The taxonomic evaluation of the 54 bacterial genomes found to harbor the IQ domain was not congruent with evolutionary ancestry but rather reflected lifestyle strategy since most of them were found to be related to known host-associated lineages. The scarce distribution (approx. 0.17% scanned genomes) mostly restricted to characterized bacterial symbionts contrasts with the much larger eukaryotic radiation and points towards HGT-mediated mobilization of IQ protein domain in Bacteria.

2. LRR_6 (PF13516) domain

- Distribution in Bacteria (GTDB R05-RS95): 4 467 sequences in 1 911 species.
- Distribution in Bacteria (Pfam 35.0 database): 1 283 sequences in 511 species.
- Distribution in Eukarya (Pfam 35.0 database): 53 469 sequences in 2 070 species.

Even though LRR_6 was found to be present in approx. 6% of bacterial species (as assessed by GTDB R05-RS95), its abundance and distribution in Eukarya show a predilection towards this domain of life. The HGT mobilization of this protein domain in Rickettsiales could not be ruled out since it was detected in 2.4% of Alphaproteobacteria genomes, the majority of which could be linked to symbiotic lineages.

3. PRANC (PF09372)

- Distribution in Bacteria (GTDB R05-RS95): 32 sequences in 5 species.
- Distribution in Bacteria (Pfam 35.0 database): 15 sequences in 3 species.
- Distribution in Eukarya (Pfam 35.0 database): 22 sequences in 7 species.
- Distribution in eukaryotic Viruses (Pfam 35.0 database): 243 sequences in 23 species.
- Distribution in prokaryotic Viruses (Pfam 35.0 database): 1 sequence in 1 species.

PRANC domain occurrences in Bacteria and Eukarya were found to be the exception rather than the norm. This very rare protein domain exhibited a narrow taxonomic distribution restricted to a few species within the Rickettsiales/ Diplorickettsiales (Bacteria) and Coleoptera/Hymenoptera/Hemiptera (Eukarya) orders. Surprisingly, it was found to be common in a variety of poxviruses (eukaryotic), where it is frequently associated with ANK repeat domains (as shown by https://pfam.xfam.org/). The phylogenetic analysis performed on PRANC domain-containing proteins showed higher sequence diversification within poxvirus clades (Fig. S11). The Rickettsiales sequences formed a monophyletic group within a larger cluster of poxvirus/eukaryotic ancestry, pointing towards PRANC domain acquisition during host co-infection. The widespread incidence of Rickettsiales among insects[14,15] together with the entomopoxvirus long-lasting host association and cytoplasmic replication capacity[16] likely increase the chances for bacteria-virus co-infections to occur.

4. PRANC (PF09372) domain in Wolbachia phage WO (GCA_002602005.1)

One PRANC domain-containing protein identified in Wolbachia phage WO displayed high sequence similarity (96-98% as assessed by blastp searches) with proteins found in Rickettsiales species belonging to the Wolbachia genus. As PRANC domain architecture is not commonly found in temperate phages (see above), its presence likely points towards a maintenance mechanism for relict (and rare) long-haul HGT events.

Long-branch extraction

The generated phylogenies pointed towards shared evolutionary history between Rickettsiales and Alpha II Rhodospirillales bacteria. This result was further scrutinized by classifying Rickettsiales proteomes (n= 128 MAGs/genomes; 171 713 proteins) using mmseqs2[6] sequence similarity searches against an Alphaproteobacteria-centric database (11 113 477 proteins belonging to 2 955 MAGs/genomes spanning 27 orders). The obtained results were further converted into a BLAST-tab formatted file (using mmseqs2 convertalis) from which individual top hits (cutoffs: E-value 1E-3, identity 10%, coverage 10%, bitscore 50) were extracted (Fig. S15).

Mitochondrial trees

An additional Bayesian mitochondria-Alphaproteobacteria phylogeny (n=33 genomes) (Fig. S16) was performed using the 24 marker genes alignment (6 416 sites alignment). The 4 chains run under the CAT-F81-dgam4 model reached convergence (maxdiff 0.04; 121 833 trees) and displayed a topology comparable to the one obtained by CAT-GTR-dgam4. Noteworthy, the CAT-F81-dgam4 phylogenomic reconstruction showed dissimilarities in i) GCA_002937725 (TMED127 lineage) and NC_021126/NC_021127 (Discoba) branching patterns. In both cases, the lineages were attracted without statistical support toward their sister groups (posterior probability values of 0.63 and 0.55, respectively) (Fig. S16). The NC_021126/NC_021127 attraction towards Ancoracysta/Opisthokonta/CRuMs branch is very likely an artifact as it broke the monophyly of Discoba. The higher number of branches with low statistical support (posterior probability values < 0.95), decreased fit in modeling site-specific amino acid propensities (z-score=3.19, in comparison with z-score=2.8 in CAT-GTR-dgam4) and Discoba monophyly breakage encouraged for CAT-GTR-dgam4 phylogeny selection.

The alignment used in Fig. 4 for generating mitochondrial-centric phylogenies (24 proteins; 6 416 aligned sites) was treated to remove highly variable sites. The site exclusion procedure was employed to evaluate if the Rickettsiales-mitochondria sister group phylogenetic relationship may be influenced by their presence. Site-stripping eliminated top 2% (depletion of approx. 3.1% parsimony-informative sites), 5% (depletion of approx. 7.7% parsimony-informative sites), and respectively 8% (depletion of approx. 12.3% parsimony-informative sites) of most heterogeneous sites present in the alignment. Obtained site-stripped alignments were subsequently used to generate phylogenies under the CAT GTR model. The phylogenies obtained with site-stripped alignments (Figs. S17-S19) showed identical topologies with the one generated using the untreated alignment. However, caution should be exercised in fast-evolving site trimming, as they may carry accurate and useful phylogenetic information[17]. Furthermore, it's very possible that zealous trimming will lead to a significant deletion of parsimony-informative sites, eliminating some of the method's key benefits and impairing phylogenetic reconstruction (due to a lack of sufficient informative sites in the alignment).

The Rickettsiales-mitochondria sister group phylogenetic relationship was further evaluated by replacing the MarineAlpha9 Bin5 (GCA_002937595) with two UBA7887 representatives (GCA_002501105 and GCA_002728255). The new genomic dataset was processed in the same way as described for the initial mitochondrial phylogeny. Briefly, the extracted sequences belonging to 24 MitoCOGs were clustered with mmseqs[6] (easy-cluster –min-seq-id 0.85 -c 0.8 –cov-mode 0). Each protein-containing cluster was independently aligned with

MAFFT-L-INS-I[18] v7.471. Hidden Markov models (HMM) were constructed from the obtained alignments using HMMER[19] v3.3 (hmmbuild). HMM-specific e-values were determined by assessing the e-values for the target (as present MitoCOGs) and non-target hits. The mitochondrial and bacterial genomes/MAGS (n=33) were screened using hmmsearch (--domblout) for the presence of the 24 MitoCOG markers. The identified proteins were independently aligned with PASTA[20] v1.12 (--estimator=raxml) and trimmed with BMGE[21] v1.12 (default settings). The trimmed alignments were further concatenated and site-stripped for the top 2% (depletion of approx. 3.1% parsimony-informative sites) and 5% (depletion of approx. 7.7% parsimony-informative sites) most heterogeneous sites. The obtained phylogenies generated through the CAT GTR model, using the untreated and site-stripped alignments (Figs. S20-S22), were found to be topologically similar to the previously obtained ones (Fig. S4; Figs. S17-S19).

Electron transport chain phylogeny

Six genes involved in oxidative phosphorylation were selected to assess the origin of the mitochondrial electron transport chain (ETC) (Table S3). The genes encoding for 'eukaryotic-type' protein complexes (as shown by KEGG: https://www.genome.jp/pathway/ko00190) and, had a mitochondrial genomic localization. The phylogenetic spread of these genes, in the prokaryotic tree of life, was evaluated through the usage of AnnoTree[22] (v1.2) (KO annotations for 27 000 bacterial and 1 500 archaeal genomes). Inquiries effectuated with the six KEGG identifiers showed a taxonomic distribution skewed towards Alphaproteobacteria. Further queries centered on the simultaneous presence of $\geq$ 3 genes decreased the number of hits and increased the observed taxonomic imbalance. Thus, the concurrent presence of the 3 NDUFS genes (Table S3) was detected in 362 genomes (out of 27 000), out of which 94% were found to be of alphaproteobacterial origin. Increasing the presence threshold to 5 genes per MAG/genome (i.e. K03878, K03935, K03941, K02256, K02132) further reduced the number of hits and constrained the taxonomic spread to Alphaproteobacteria (42 genomes, 85.7% belonging to Rickettsiales).

Phylogeny-aware multiple sequence alignments (MSAs) were constructed for each phylogenomic marker (n=6) using the software PASTA[20] v1.8.3 (Mirarab et al., 2015) with default settings. The obtained MSAs were concatenated in a supermatrix (2 493 aligned sites) before performing ETC-focused phylogenies. 4 independent chains were run through PhyloBayes MPI[23] 1.8b with: i) site-specific equilibrium frequency profiles (-cat option), ii) uniform exchangeabilities (-f81 option) and iii) rate variation across sites modeled through a discretized gamma distribution with four categories (-dgam 4). The chains were stopped after approx. 121 000 trees (for each chain) and their burn-in (i.e., the number of points before the chain has reached stationarity) estimated at 13 500 points by Tracer[24] v1.7.1. The convergence of parameters and tree space between the 4 chains was assessed by tracecomp and bpcomp software (as implemented in PhyloBayes MPI 1.8b). The obtained values for effective sizes (> 27 000), discrepancies (< 0.05), and the maximum difference in bipartition frequencies (= 0.02) indicated convergence. Thus, a consensus tree was calculated by pooling the trees (approx. 450 000) from the 4 chains.

Despite their minute (in content) and simplified genomes, mitochondria commonly retained genes involved in translation and ATP generation. 6 genes involved in oxidative phosphorylation were selected to reconstruct the evolutionary history of the ETC. The distribution of these genes within the prokaryotic tree of life revealed a narrow taxonomic range, with a skew towards several Alphaproteobacteria radiations. Among these lineages, Rickettsiales members were found to harbor the highest numbers per genome. The generated Bayesian phylogeny pictured a topology in which mitochondria and basal Rickettsiales displayed high support (posterior probability = 0.95) sister-group relationship. Posterior predictive analyses effectuated on model-generated data (CAT-F81) and the original sequence alignment showed that the evolutionary reconstruction adequately modeled the observed site-specific amino acid propensities (z-score=0.51, p=0.3). The close phylogenetic connection between the ETCs of Rickettsiales and mitochondria implies that they speciated from a common evolutionary lineage, which had already acquired the ability to perform aerobic respiration. The observed pattern of ancestry and descent depicts basal Rickettsiales as the closest phylogenetic lineage to mitochondria (to date) and suggests that the mitochondrial ancestral radiation thrived in a (micro)oxic environment. This inference is further corroborated by the presence of oxygen-dependent pathways in all Rickettsiales families.

Nuclear-encoded mitochondrial/mitochondrial homologous genes present in Rickettsiales genomes

Despite their gene-impoverished genomes, mitochondria (in general) are solely responsible for powering the cell's intricate metabolic circuitries[25]. Their minute (in content) and simplified genomes of proteobacterial ancestry commonly retained genes involved in translation and ATP generation[25]. Additional genes encoding for ribosomal proteins, RNA processing, and protein import/maturation are sometimes also detected, but their presence is associated with a restricted phylogenetic distribution within Eukarya[26]. While many of the mitochondrial genes have been lost, some of them have been transferred to the nuclear genome from where their products are imported back into the organelle where they function.

A. Oxygen-dependent ubiquinone biosynthetic pathway

- **K06127**: COQ5- 2-methoxy-6-polyprenyl-1,4-benzoquinol methylase; **K06134**: COQ7- 3-demethoxyubiquinol 3-hydroxylase; **K18587**: COQ9- ubiquinone biosynthesis protein COQ9; **K18588**: COQ10- coenzyme Q-binding protein COQ10
- Distribution in Bacteria (GTDB R05-RS95): K06127 + K06134 + K18587 + K18588; **946** hits (100% Alphaproteobacteria). Absent in Archaea.
- Isoprenoid quinones are membrane-bound molecules (comprising a polar quinone ring and a hydrophobic isoprenoid side chain) present within the membranes of most living organisms. Here, they perform key bioenergetic functions by partaking in the electron/proton transfers in the respiratory electron transport chain. The large majority of naturally occurring isoprenoid quinones belong to either naphthoquinones (e.g., menaquinones), or the evolutionary younger benzoquinones (e.g., rhodoquinones, plastoquinones, and ubiquinones). While menaquinones are present in most bacterial and archaeal phyla and considered to be 'as old' as the last universal common ancestor, ubiquinones are taxonomically restricted to several proteobacterial and eukaryotic (where they participate in mitochondrial oxidative phosphorylation) lineages[27]. While it is likely that the eukaryotic ubiquinone pathway was inherited from the mitochondrial progenitor its evolutionary history is not well understood. The oxygen-dependent ubiquinone biosynthetic pathway in eukaryotes and most Proteobacteria diverges at the level of 4-Hydroxy-3-polyprenylbenzoate, to converge later at 2-Polyprenyl-6-methoxyphenol (see https://www.genome.jp/pathway/map00130). The last three steps of the metabolic pathway are performed by catalytically equivalent but structurally unrelated nuclear-

encoded enzymes (i.e., COQ5/COQ7/COQ3 in Eukarya and UbiE/UbiF/UbiG in Proteobacteria).

B. Aerobic electron transport chain

- **K03937**: NADH dehydrogenase (ubiquinone) Fe-S protein 4 (NDUFS4); **K00234**: succinate dehydrogenase (ubiquinone) flavoprotein subunit (SDHA SDH1).
- Distribution in Bacteria (GTDB R05-RS95): K03937 + K00234; **89** hits (100% Alphaproteobacteria). Absent in Archaea.
- Important elements of the aerobic mitochondrial electron transport chain include the nuclear-encoded subunit 1 (SDHA) of succinate dehydrogenase and the mitochondrial-encoded subunit 4 of NADH dehydrogenase (NDUFS4). Both NDUFS4 and SDHA are found in the inner mitochondrial membrane (in eukaryotes) where they play a role in electron transfer. Both these proteins are thought to have been present in the mitochondrial ancestor[28].

C. Miscellaneous

- **K19054**: frataxin (FXN); **PF06258**: mitochondrial fission ELM1.
- Distribution in Bacteria (GTDB R05-RS95): K19054 + PF06258; **99** hits (100% Alphaproteobacteria). Absent in Archaea.
- Frataxin is produced in the cytoplasm before being transported into the mitochondria, like the majority of nuclear-encoded mitochondrial proteins. Though the exact function of frataxin is debatable, it has been shown that this protein is involved in various processes ranging from iron homeostasis and oxidative phosphorylation to heme metabolism and protection against oxidative damage[29,30]. The other nuclear-encoded mitochondrial protein ELM1 was described to participate in mitochondrial fission in plants, where it plays a role in moving dynamin-related proteins from the cytosol to mitochondrial fission sites[31]. Although this protein has previously been found in bacteria, its functionality remains unknown.

These 8 genes (in eukaryotes seven nuclear and one mitochondrial encoded) that were inferred to have been present in the pre-mitochondrial ancestor[28] were simultaneously present in 37 genomes (out approx. 31 910). This limited taxonomic distribution within the bacterial line of descent (exclusively restricted to Alphaproteobacteria), together with their presence in ancient Rickettsiales lineages (six genomes belonging to ancient stage Rickettsiales and one to UBA1997) points towards common evolutionary history. While gene distribution patterns are not sufficient for inferring ancestry and descent, they strengthen the Rickettsiales-mitochondria sister-group relationship obtained by evolutionary history reconstructions.

<u>5-Aminolevulinic acid (ALA) and carnitine/acylcarnitine transporter (CACT) phylogenies</u>

5-Aminolevulinic acid (ALA) is a non-proteinogenic amino acid that acts as a precursor to all tetrapyrrole compounds, including chlorophyll, heme, and vitamin $B_{12}$. Tetrapyrrole biosynthesis departure point is represented by ALA that can be generated through either C4 (Shemin pathway) or C5 biosynthetic pathways. While C4 was found to be restricted to several eukaryotic (mammals, fungi, and protists) and prokaryotic (Alphaproteobacteria) lineages, C5 was determined to occur in most Bacteria, Archaea and plastid-containing eukaryotes[32] (for an exception see here Foley et al.[33]). In the C4 pathway ALA is produced in a single step by 5-Aminolevulinate synthase (ALAS) that catalyzes the condensation of succinyl-CoA and glycine. In eukaryotes, the nuclear-encoded ALAS is translated into a precursor protein that is further directed by an N-terminal signal sequence into the inner mitochondrial matrix. Here, following signal sequence removal the enzyme makes use of the available glycine and citric acid cycle intermediate succinyl-CoA[34]. The existence of two ALA biosynthetic pathways within eukaryotes was previously interpreted as the result of endosymbiotic gene transfers (from mitochondria and chloroplast, respectively) followed by the selective retention of the C5 pathway in plastid-containing organisms[32].

Performed AnnoTree (v1.2) searches with K00643 and TIGR01821 identifiers revealed the absence of ALAS-encoding genes in Archaea and their skewed taxonomic distribution within Bacteria. In the latter, most of the occurrences were found to be restricted to members of the Alphaproteobacteria class (85.5% of K00643 and 93.5% of TIGR01821 hits). While a phylogenetic affinity between basal clades of ancient stage Rickettsiales (SFRX01, UBA3002, and UBA998) and eukaryotes can be observed in ALAS phylogeny (Fig. S23), the low statistical support for this monophyletic clade prevents confident evolutionary ancestry inferences. Low branch support values are likely a consequence of reduced sequence conservation (average identity=48%) and alignment length.

The very restricted taxonomic spread of the CACT gene (20 genome hits for K15109 in AnnoTree v1.2) within the bacterial line of descent, together with its presence in known bacterial symbiotic lineages (i.e., Legionella; Paracaedimonas; Neochlamydia; Tatlokia) points towards acquisition through horizontal gene transfers. The occurrence of CACT in the late ancient Rickettsiales UBA6187 family suggests a likely eukaryotic association, which probably set the stage for the emergence of transition/intracellular lineages.

CARD-FISH probe used for cryptophytes identification

- Probe name: Crypto B[35,36]
- Target group: Cryptophyceae
- Coverage: 74.3%
- Sequence 5’-3’: ACGGCCCCAACTGTCCCT
- Formamide concentration (%): 50

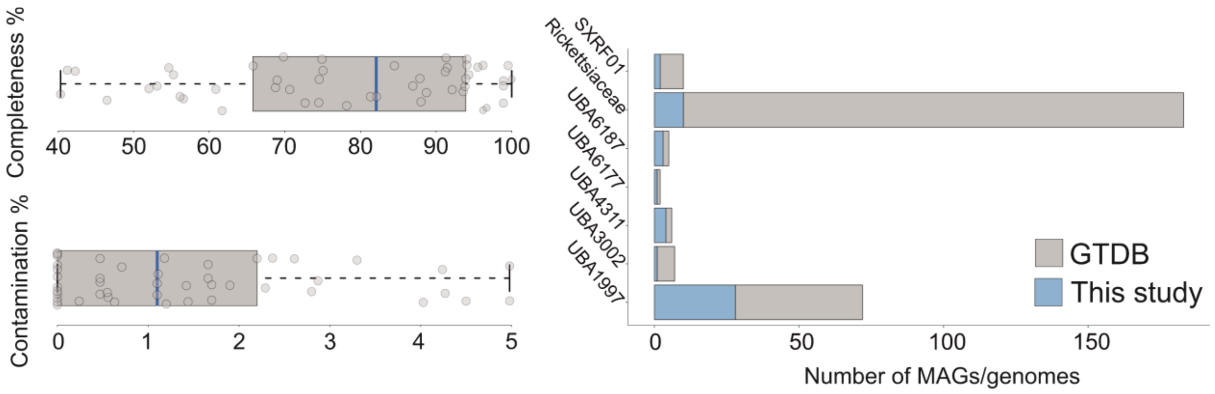


Fig. S1. **Qualitative and taxonomic assessments of the recovered Rickettsiales MAGs (n=49)**. The left panel of the figure depicts the estimated levels of completeness and contamination, while the right one shows the family-level classification in relation to publicly available Rickettsiales genomes/MAGs.

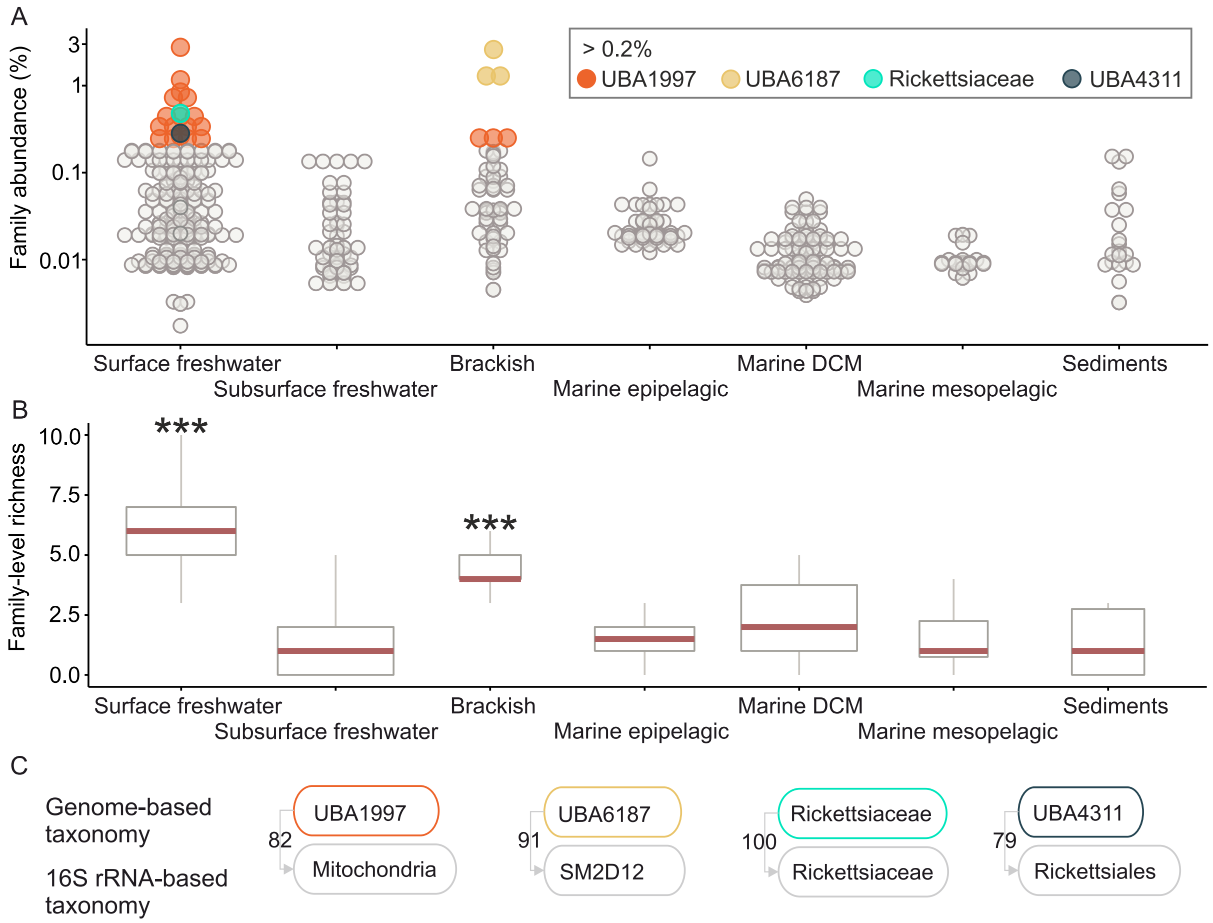


Fig. S2. **Rickettsiales diversity in aquatic ecosystems**. (A) The family-level abundance of Rickettsiales bacteria in seven habitats. The X-axis shows the metagenomic datasets (n=212) grouped by environment type; the Y-axis (log10 scale) indicates the percentages of each family-level group (n=12) within the prokaryotic communities. The figure's inset (upper right panel) depicts the taxonomy of Rickettsiales families with abundances higher than 0.2%. (B) Family-level richness across seven habitat types. The X-axis shows the metagenomic datasets grouped by the environment; the Y-axis indicates their taxonomic richness. Black-colored stars highlight habitats with significantly higher richness (Kruskal-Wallis, p-value < 2.2e-16). DCM: deep chlorophyll maxima. (C) The figure shows the taxonomic equivalence (in % identity) between genomic-based taxonomy and the 16S rRNA gene-based one (as assessed by 16S rRNA genes classification) for the Rickettsiales families with abundances higher than 0.2%.

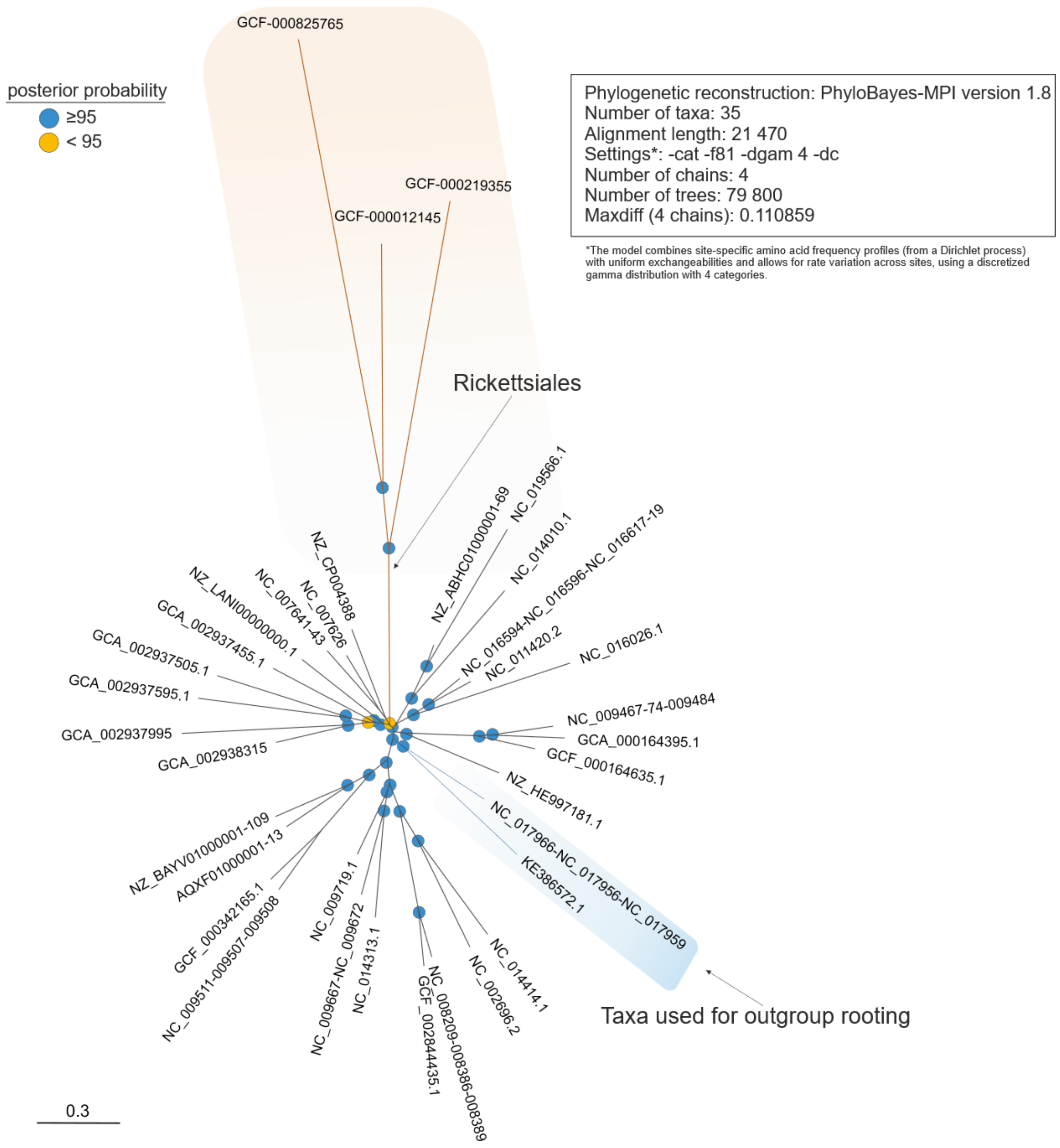


Fig. S3. **Unrooted Bayesian phylogeny (-cat -f81 -dgam4) of slow-evolving Alphaproteobacteria lineages and selected Rickettsiales genomes**. The blue-colored box highlights the branch selected as an outgroup for downstream Rickettsiales phylogenetic reconstructions. Outgroup choice was driven by: i) branch positioning (cladistically outside Rickettsiales), ii) phylogenetic proximity, and iii) the presence of slow-evolving taxa. Posterior probability values are represented through colored circles (top left legend). The scale bar indicates the number of substitutions per site.

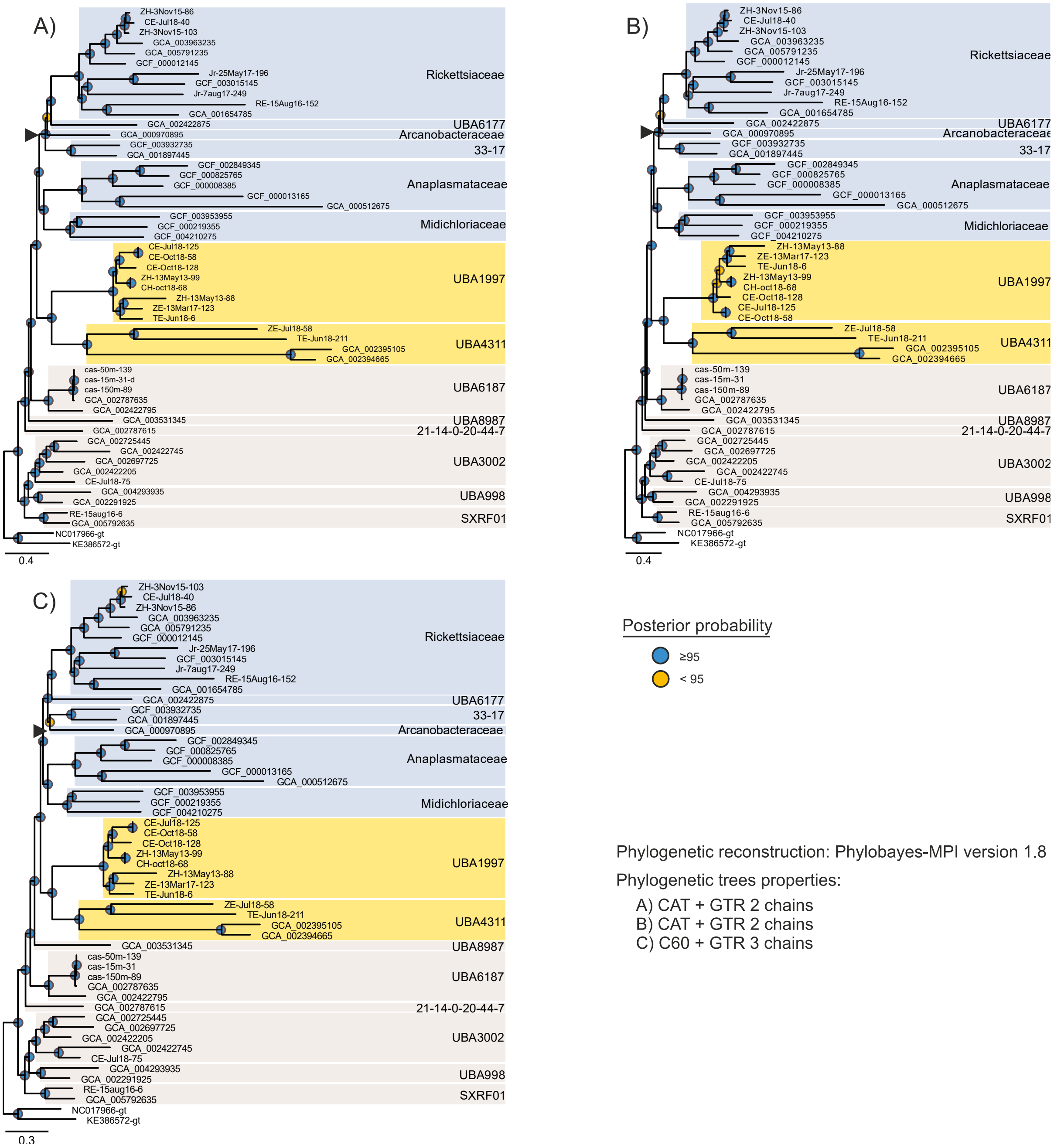


Fig. S4. **Rickettsiales Bayesian phylogenies generated under CAT+GTR and C60+GTR evolutionary models**. The black-colored triangle highlights the Arcanobacteraceae lineage. Posterior probability values are represented through colored circles (right legend). The scale bar indicates the number of substitutions per site.

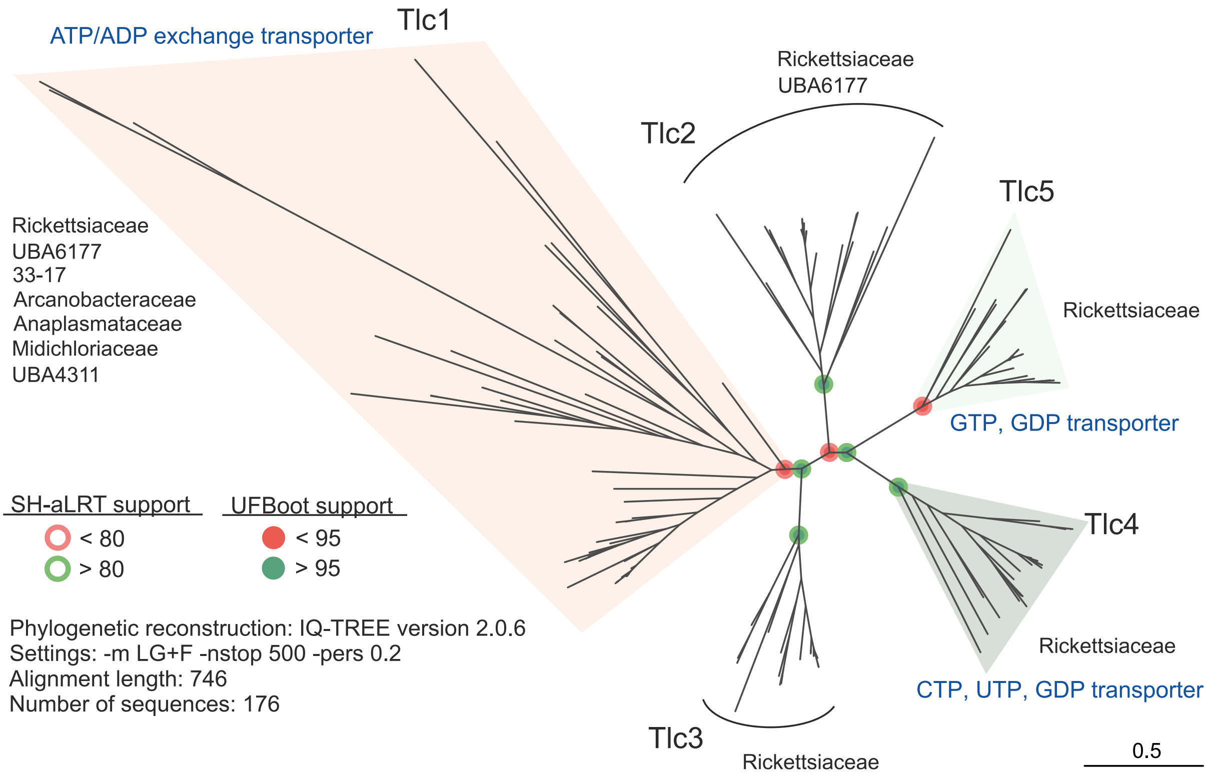


Fig. S5. **Maximum-likelihood phylogeny (LG+F, general amino acid exchange rate matrix with empirical base frequencies) of Rickettsiales ATP/ADP translocases**. Phylogeny-based functional inferences were built on the study of Audia and Winkler[37]. Colored boxes highlight translocases clades (Tlc1, Tlc4, and Tlc5) with known functionality (see blue-colored annotations). Family-level taxonomic composition is depicted for each clade (Tlc1-Tlc5). Branch support values (UFBoot and Sh-aLRT) are indicated through colored circles (see the legend on the left side of the figure). The scale bar indicates the number of substitutions per site.

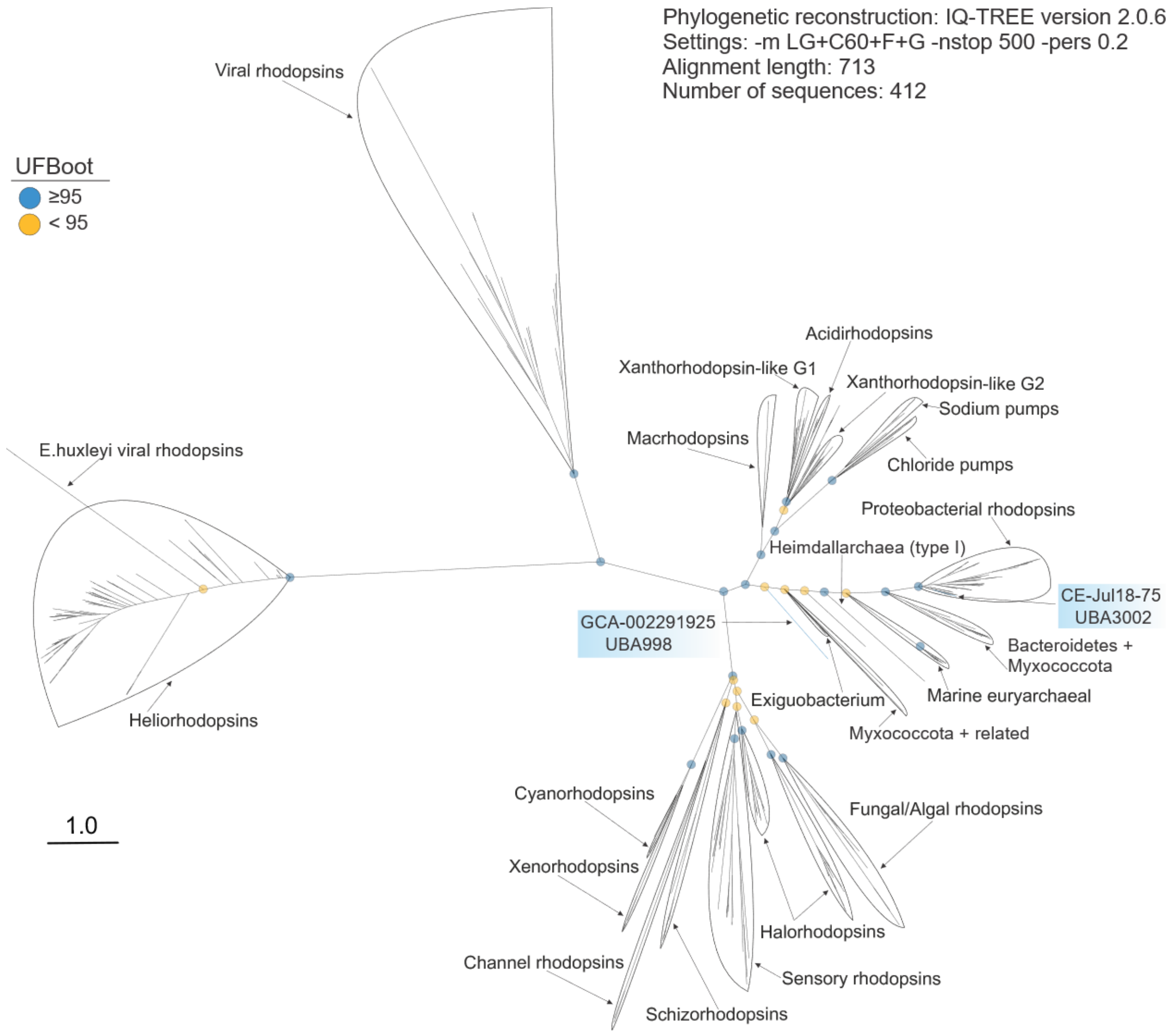


Fig. S6. **Rhodopsins maximum-likelihood phylogeny (LG+C60+F+G, general amino acid exchange rate matrix with 60-profile mixture models, empirical base frequencies, and Gamma rate heterogeneity)**. Rhodopsin reference sequences (as well as functional and taxonomic annotations) were recovered from Bulzu, Andrei, et al.[38]. Blue-colored boxes highlight Rickettsiales rhodopsins. Branch support values (UFBoot) are indicated through colored circles (see the legend on the left side of the figure). The scale bar indicates the number of substitutions per site.

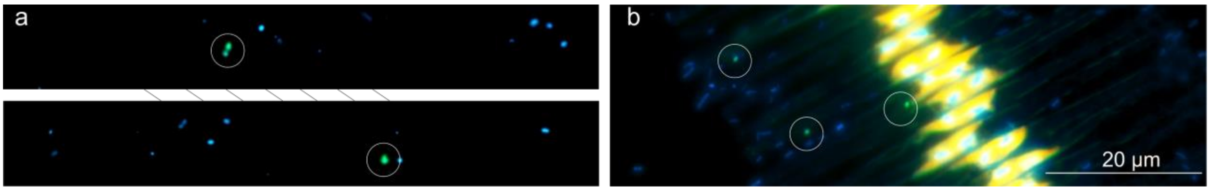


Fig. S7. **Environmental-derived microphotographs of UBA3002 Rickettsiales**. Family members are pelagic (a) or present on the surface of a Fragilaria diatom (b). Both images represent three channel overlays (DAPI, FITC, and chlorophyll). The scale bar size corresponds to 20 µm.

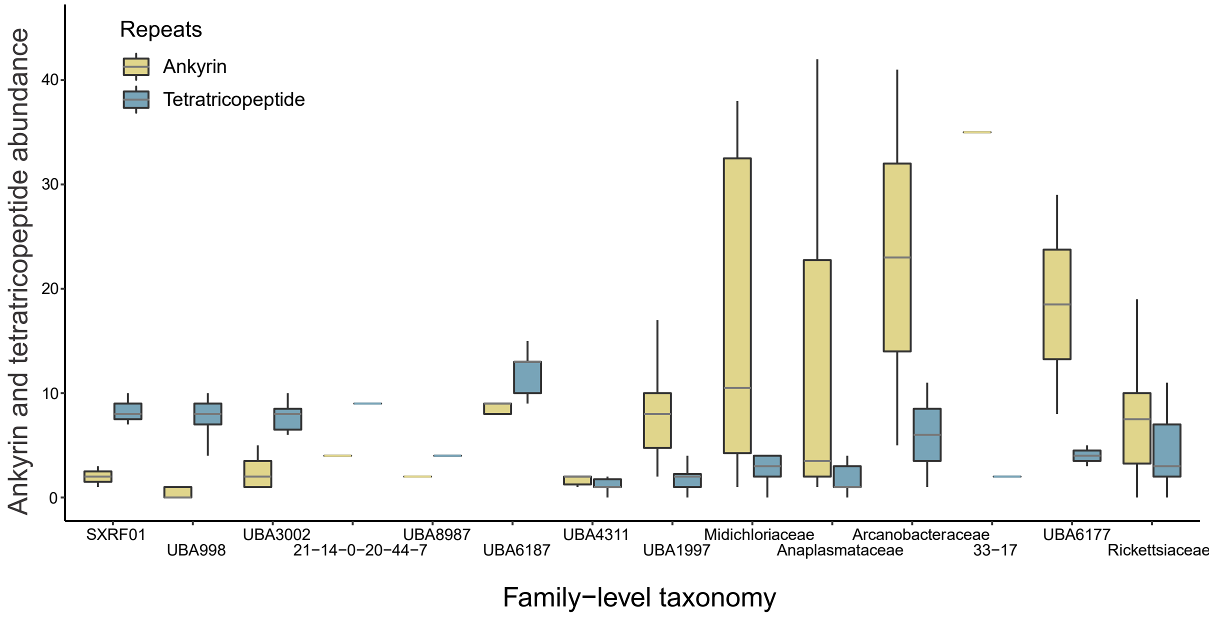


Fig. S8. **Protein scaffold abundance**. The figure depicts the abundance of ankyrin (yellow) and tetratricopeptide (blue) scaffolds in Rickettsiales families.

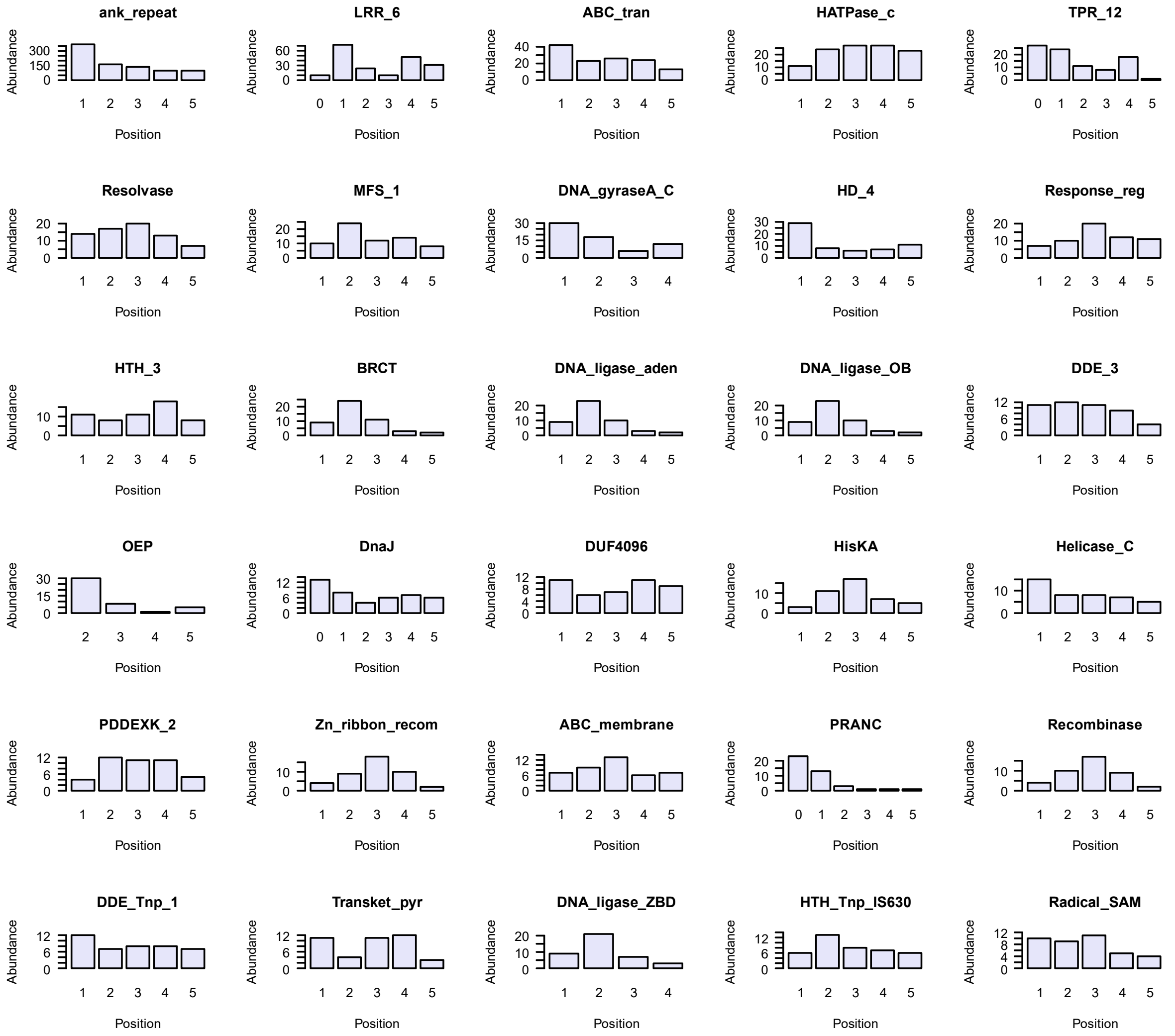


Fig. S9. **Ankyrin gene-context analysis**. The figure shows the abundance of selected top 30 most abundant co-occurring protein domains (± 5 proteins situated upstream and downstream of an ankyrin-containing one).

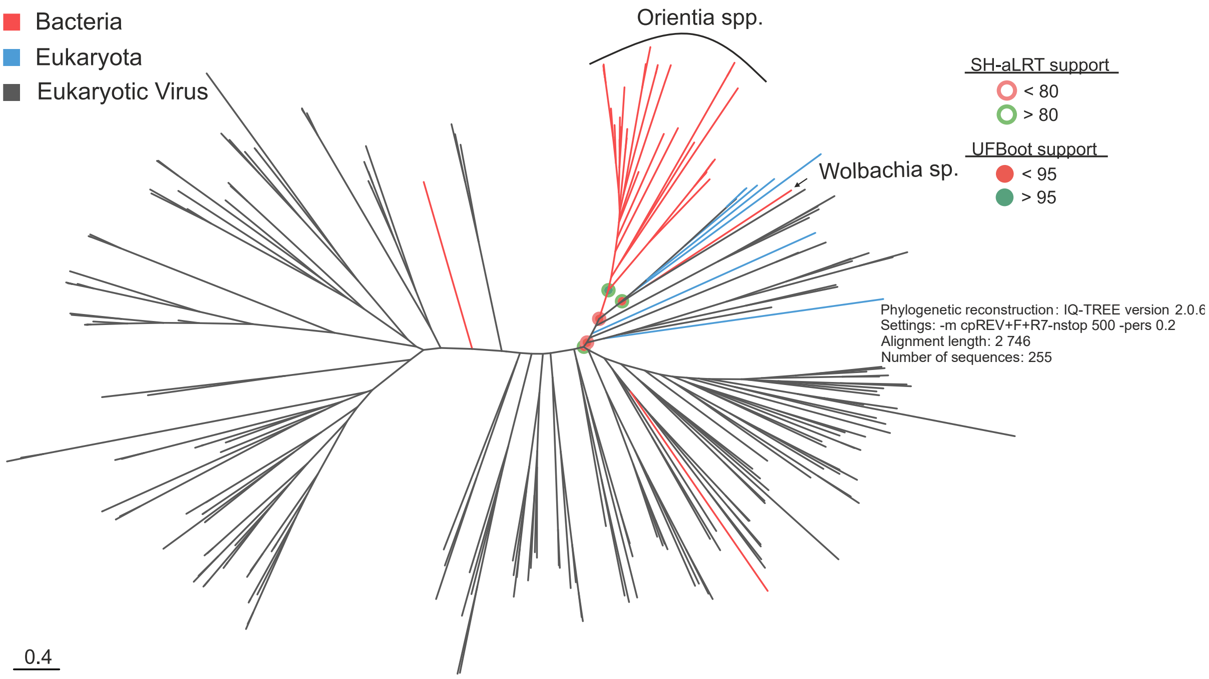


Fig. S10. **Maximum-likelihood phylogeny of PRANC domain-comprising proteins**. Branch support values (UFBoot and Sh-aLRT) are indicated through colored circles (see the legend on the right side of the figure). The scale bar indicates the number of substitutions per site.

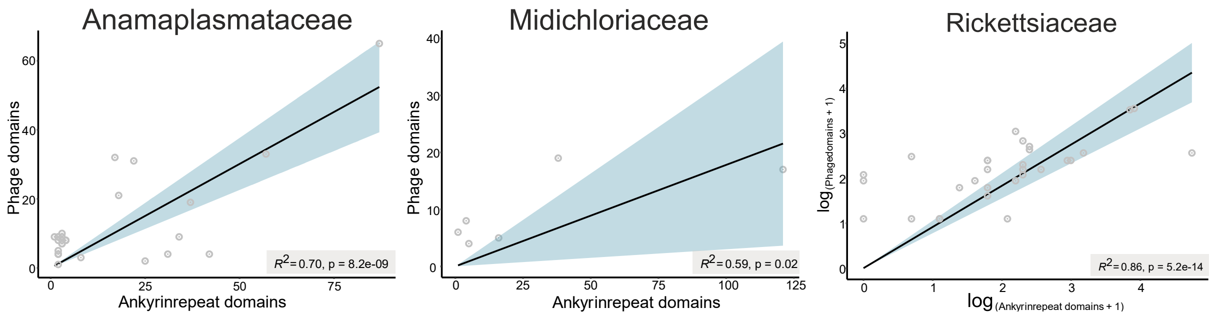


Fig. S11. **Correlations between phage and ankyrin protein domains**. The Y-axis represents the number of phages found per genome and the X-axis depicts the number of Ankyrins per genome. The figure is separated into three plots corresponding respectively to Rickettsiales families: i) Anaplasmataceae (left), ii) Midichloriaceae (middle) and iii) Rickettsiaceae (right). The coefficient of determination (R-squared) was found to be significant among the three correlations.

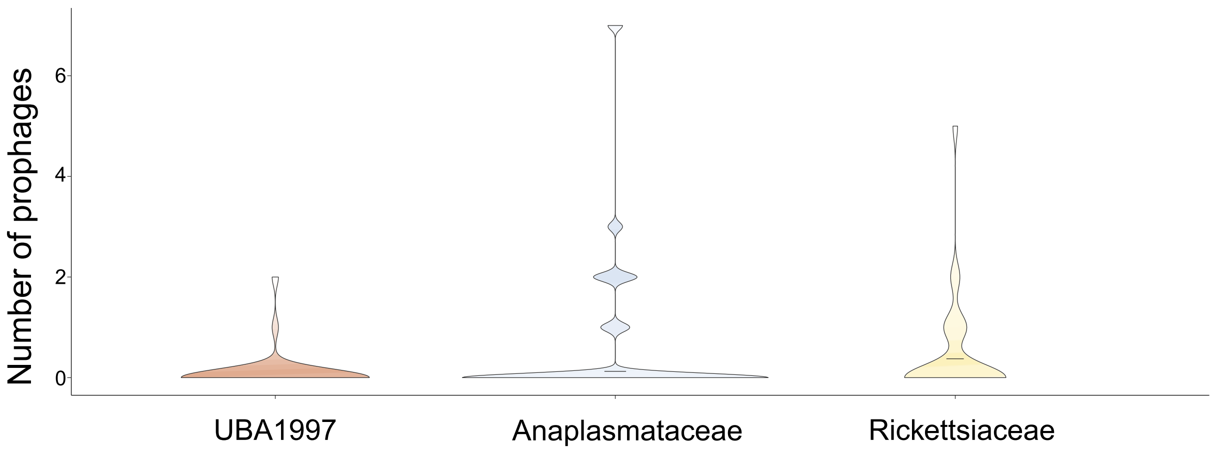


Fig. S12. **Prophages distribution in selected Rickettsiales lineages**. The figure highlights the higher number of prophages harbored by the intracellular lineages (albeit with no statistical support). Transition (UBA1997 n=32) and intracellular stage families (Anaplasmataceae n=28; Ricketssiaceae n=30) were selected based on their high number of available genomes.

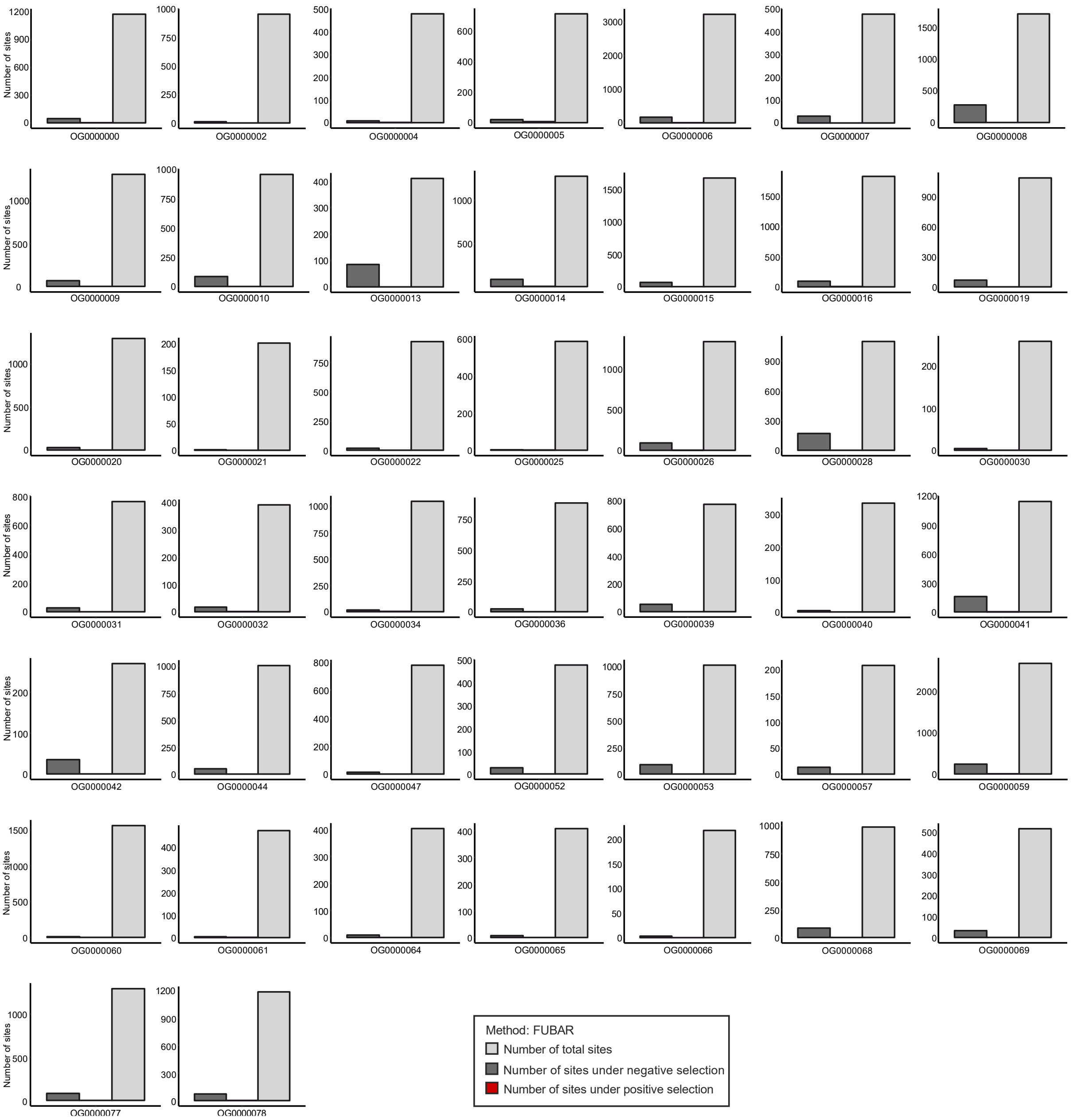


Fig. S13. **Selection pressure acting on transposases harbored by intracellular Rickettsiales families (Midichloriaceae, Anaplasmataceae, Arcanobacteriaceae, and Rickettsiaceae)**. The Y-axis highlights the number of sites under negative (purifying) or positive (diversifying) selection in 44 transposases orthologous groups (n=969 sequences). The X-axis designates individual orthologous group identifiers.

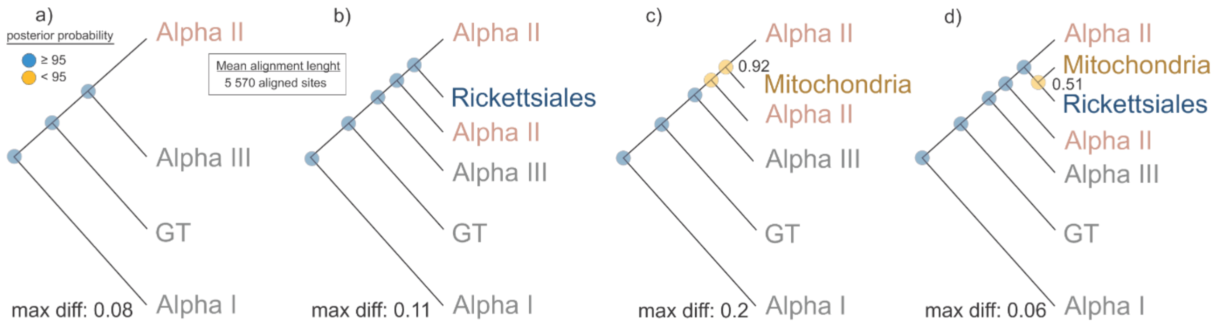


Fig. S14. **Schematic representation of long-branch extraction approach applied to Rickettsiales and mitochondrial fast-evolving lineages**. a) Backbone phylogeny of Alphaproteobacteria constructed using slow-evolving lineages. b) Backbone phylogeny of Alphaproteobacteria together with the fast-evolving Rickettsiales lineage. C) Backbone phylogeny of Alphaproteobacteria together with the fast-evolving mitochondrial lineage. D) Backbone phylogeny of Alphaproteobacteria together with the fast-evolving Rickettsiales and mitochondrial lineages. All consensus phylogenies (n chains=4) were generated through Bayesian inference (CAT+GTR). Posterior probability values are represented through colored circles (top left legend). Posterior probability values are shown between mitochondria and its closest phylogenetic lineage.

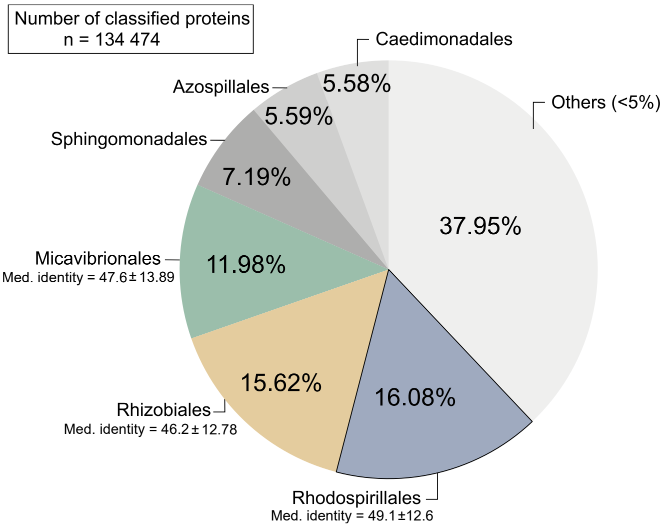


Fig. S15. **Order-level taxonomic classification of Rickettsiales proteomes based on top hits similarity search results**. The pie chart highlights the percentages of proteomes associated with a particular Alphaproteobacterial order. Taxonomic categories with hits lower than 5% were amassed in the artificial category ‘others’.

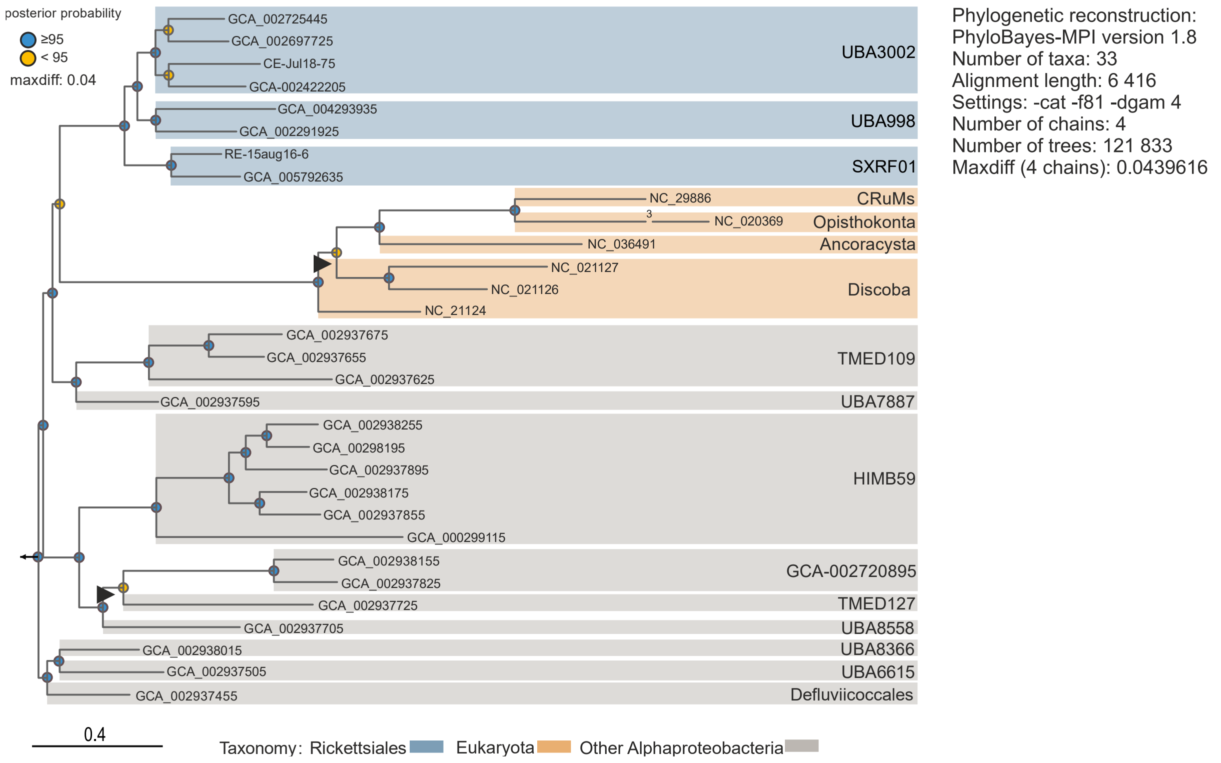


Fig. S16**. Mitochondrial-centric genome-based phylogeny generated through Bayesian inference (-cat -f81 -dgam4)**. Posterior probability values are represented through colored circles (top left legend). Taxonomical affiliation is highlighted through the usage of colored panels (see bottom legend). Black triangles highlight branching pattern discrepancies (in contrast to the -cat -gtr -dgam4 tree presented in Fig. 4a). The length of the fast-evolving *Nuclearia simplex* mitochondrial branch (NC_020369) is shown reduced (3× the scale bar). The scale bar indicates the number of substitutions per site.

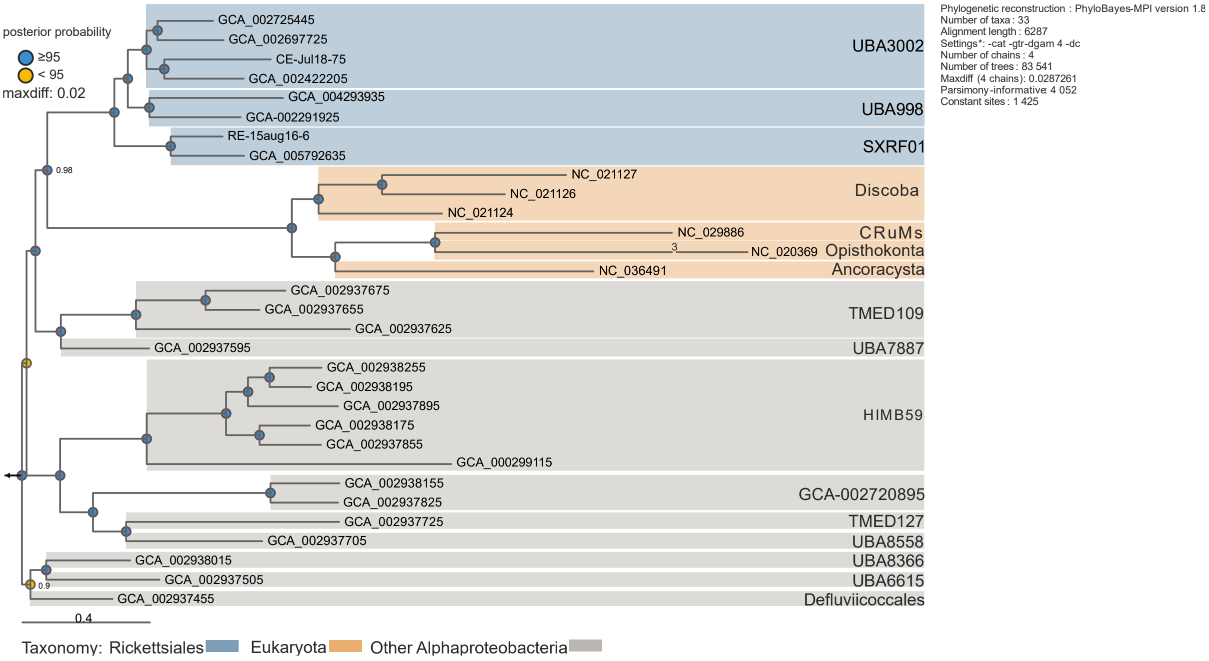


Fig. S17. **Mitochondrial-centric genome-based phylogeny generated through Bayesian inference (-cat -gtr -dc)**. The top 2% of most heterogeneous sites were removed from the alignment. Posterior probability values are represented through colored circles (top left legend). Taxonomical affiliation is highlighted through the usage of colored panels (see bottom legend). The length of the fast-evolving *Nuclearia simplex* mitochondrial branch (NC_020369) is shown reduced (3× the scale bar). The scale bar indicates the number of substitutions per site.

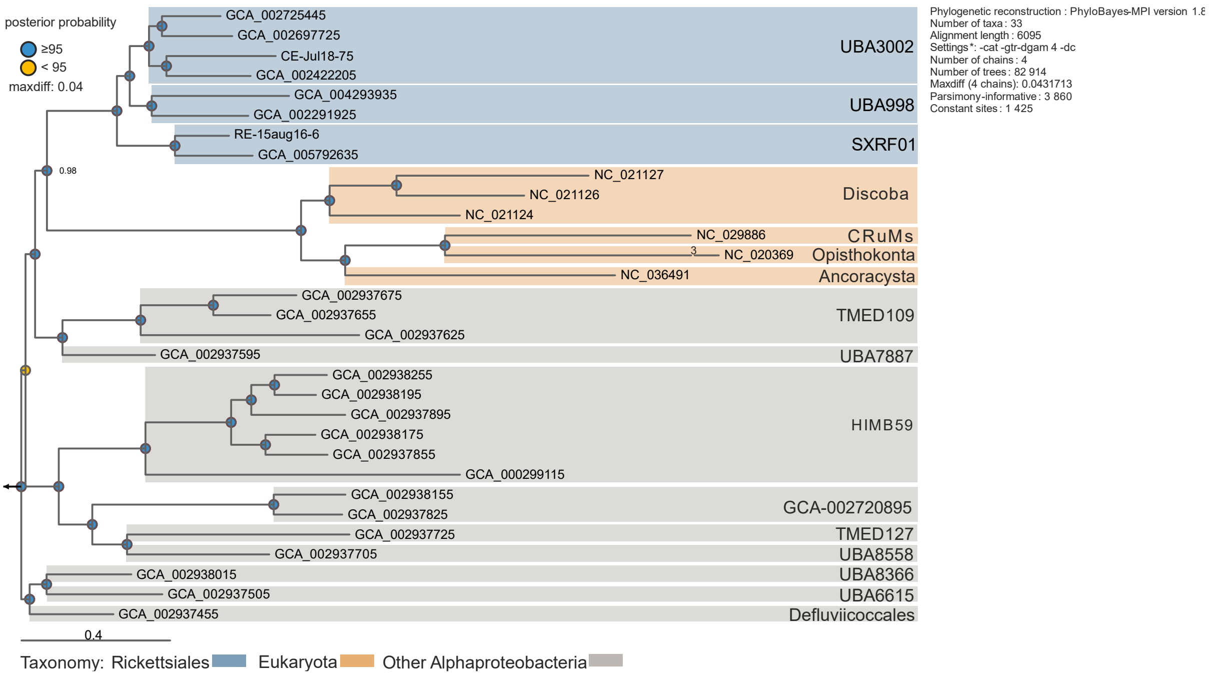


Fig. S18. **Mitochondrial-centric genome-based phylogeny generated through Bayesian inference (-cat -gtr -dc)**. The top 5% of most heterogeneous sites were removed from the alignment. Posterior probability values are represented through colored circles (top left legend). Taxonomical affiliation is highlighted through the usage of colored panels (see bottom legend). The length of the fast-evolving *Nuclearia simplex* mitochondrial branch (NC_020369) is shown reduced (3× the scale bar). The scale bar indicates the number of substitutions per site.

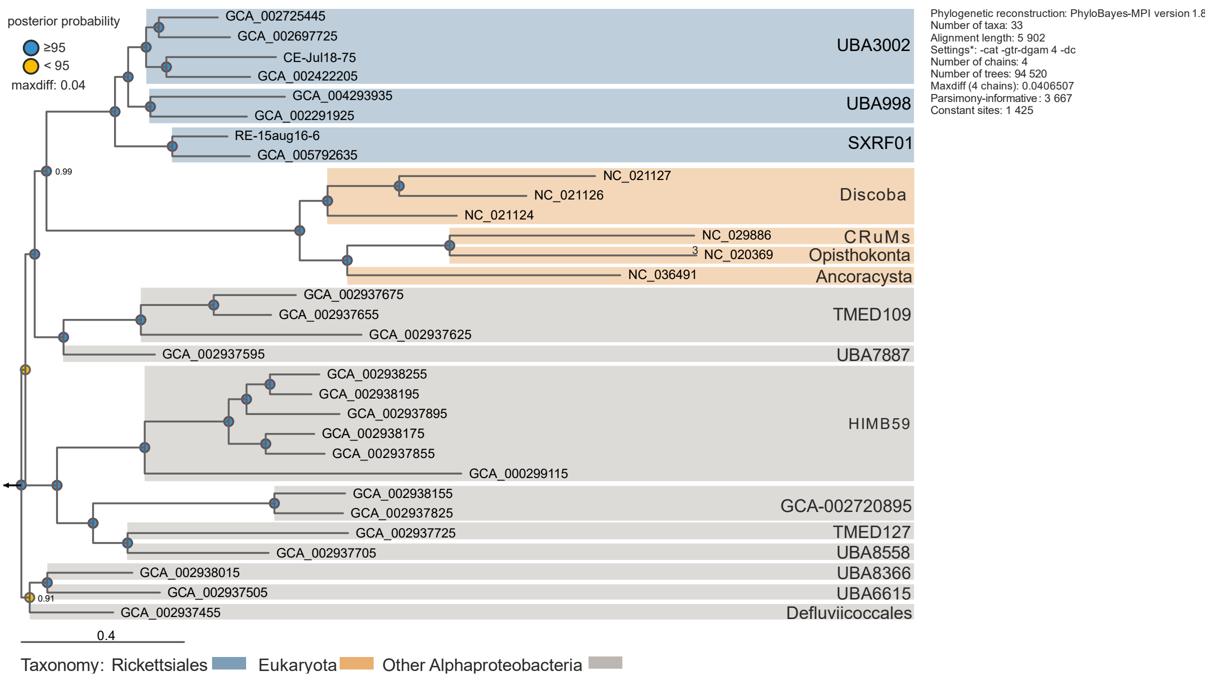


Fig. S19. **Mitochondrial-centric genome-based phylogeny generated through Bayesian inference (-cat -gtr -dc)**. The top 8% of most heterogeneous sites were removed from the alignment. Posterior probability values are represented through colored circles (top left legend). Taxonomical affiliation is highlighted through the usage of colored panels (see bottom legend). The length of the fast-evolving *Nuclearia simplex* mitochondrial branch (NC_020369) is shown reduced (3× the scale bar). The scale bar indicates the number of substitutions per site.

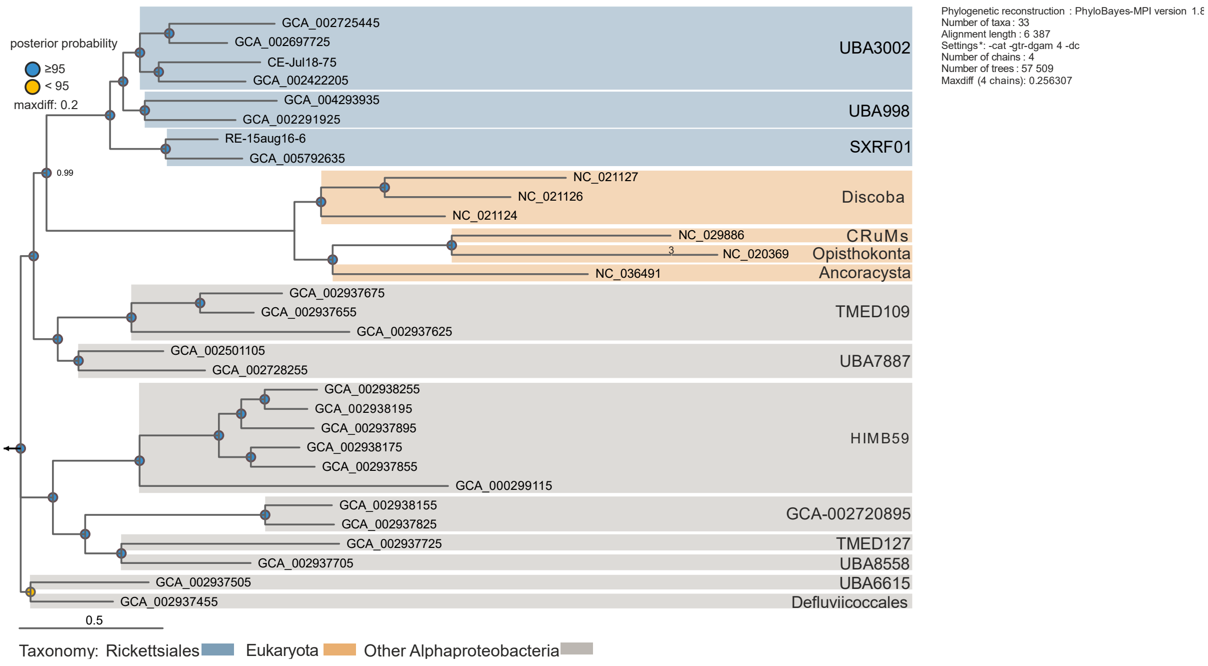


Fig. S20. **Mitochondrial-centric genome-based phylogeny generated through Bayesian inference (-cat -gtr -dc) with MarineAlpha9 Bin5 replacement (GCA_002937595).** Posterior probability values are represented through colored circles (top left legend). Taxonomical affiliation is highlighted through the usage of colored panels (see bottom legend). The length of the fast-evolving *Nuclearia simplex* mitochondrial branch (NC_020369) is shown reduced (3× the scale bar). The scale bar indicates the number of substitutions per site.

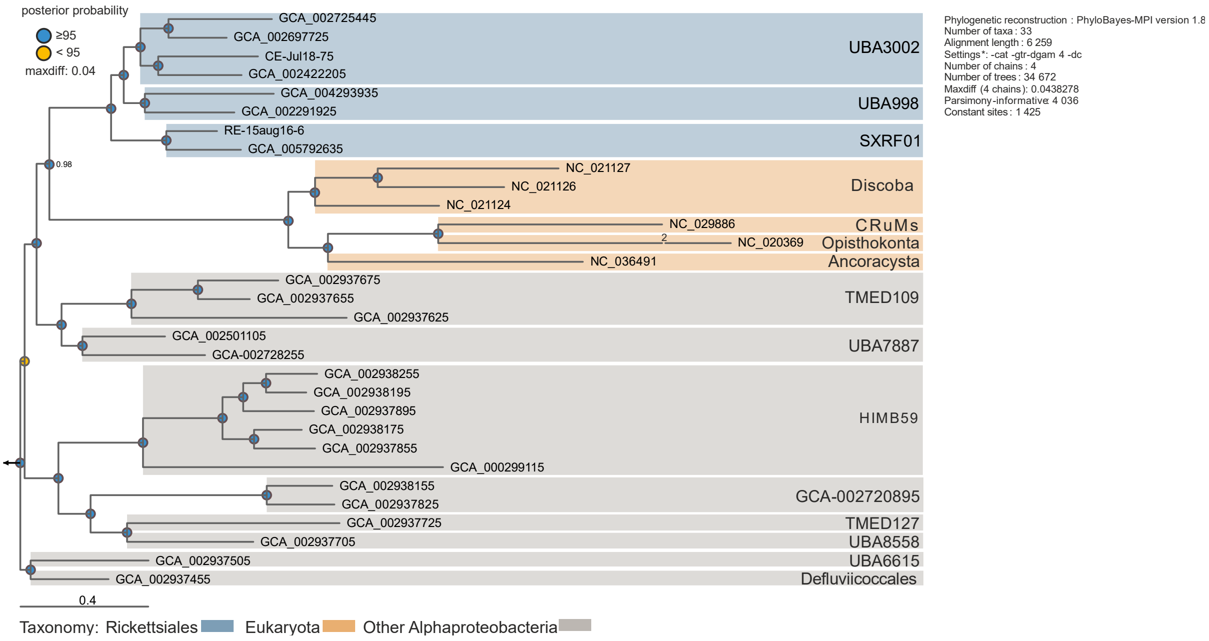


Fig. S21. **Mitochondrial-centric genome-based phylogeny generated through Bayesian inference (-cat -gtr -dc) with MarineAlpha9 Bin5 replacement (GCA_002937595)**. The top 2% of most heterogeneous sites were removed from the alignment. Posterior probability values are represented through colored circles (top left legend). Taxonomical affiliation is highlighted through the usage of colored panels (see bottom legend). The length of the fast-evolving *Nuclearia simplex* mitochondrial branch (NC_020369) is shown reduced (3× the scale bar). The scale bar indicates the number of substitutions per site.

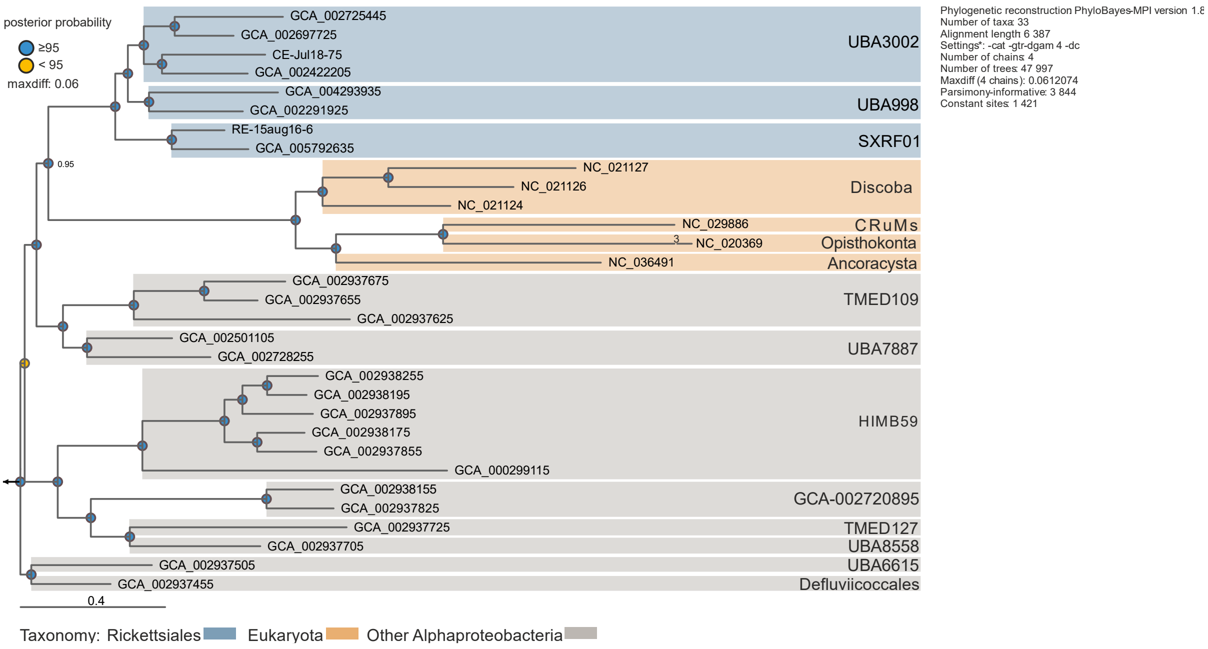


Fig. S22. **Mitochondrial-centric genome-based phylogeny generated through Bayesian inference (-cat -gtr -dc) with MarineAlpha9 Bin5 replacement (GCA_002937595)**. The top 5% of most heterogeneous sites were removed from the alignment. Posterior probability values are represented through colored circles (top left legend). Taxonomical affiliation is highlighted through the usage of colored panels (see bottom legend). The length of the fast-evolving *Nuclearia simplex* mitochondrial branch (NC_020369) is shown reduced (3× the scale bar). The scale bar indicates the number of substitutions per site.

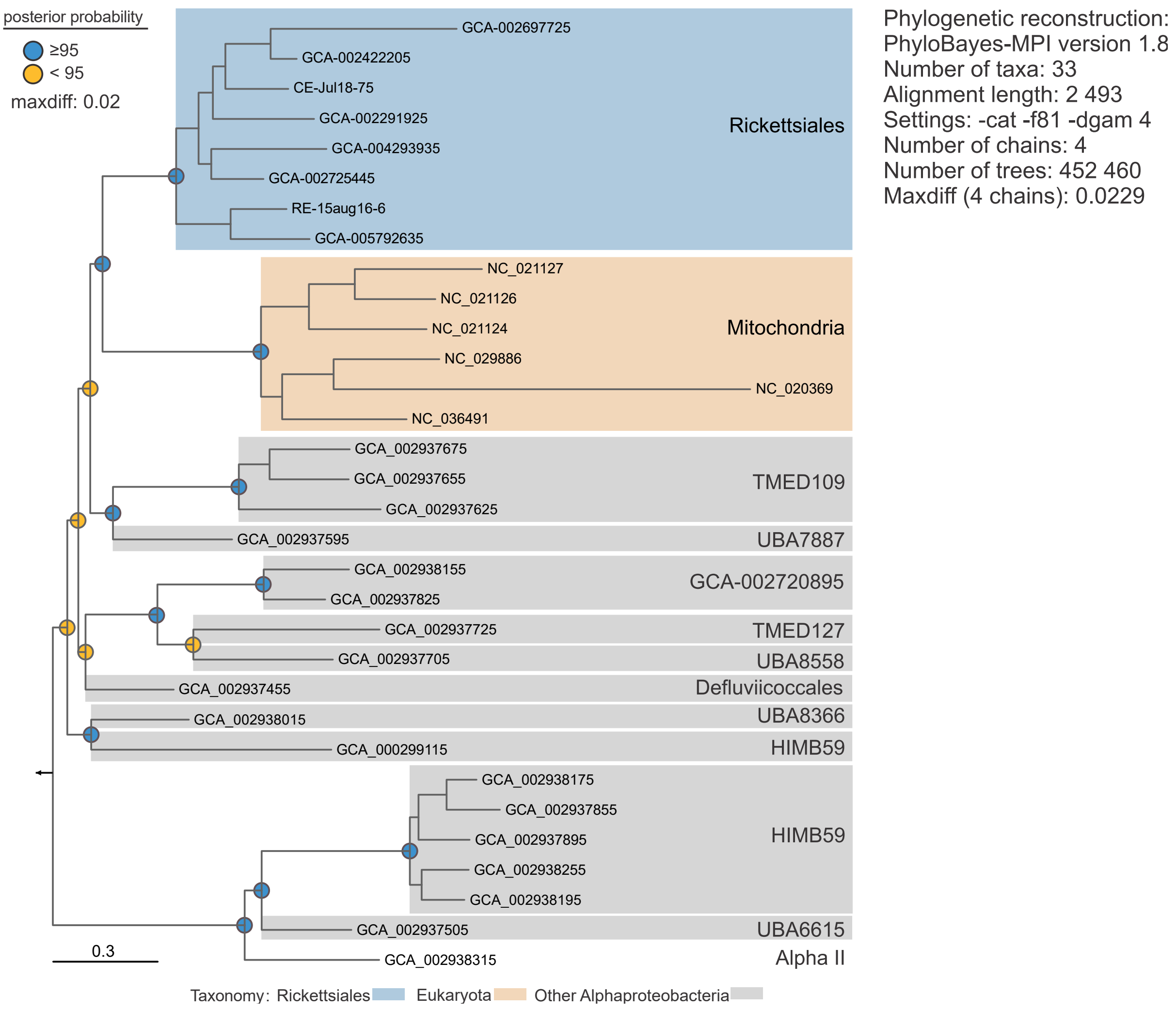


Fig. S23. **Electron transport chain phylogeny generated through Bayesian inference (-cat -f81 -dgam4)**. Posterior probability values are represented through colored circles (top left legend). Taxonomical affiliation is highlighted through the usage of colored panels (see bottom legend). The scale bar indicates the number of substitutions per site.

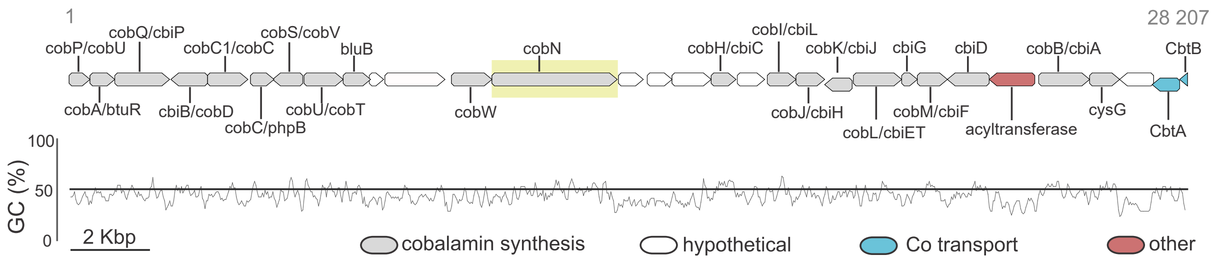


Fig. S24. **Adenosylcobalamin biosynthesis gene cluster in SXRF01 (ancient stage Rickettsiales; ss-metabat2.31 MAG)**. Genes predicted to be involved in *de novo* adenosylcobalamin synthesis are depicted in grey color. A yellow panel highlights the cobN gene- indicator for the aerobic biosynthetic pathway (cobS and cobT are also present but not part of the gene cluster). The scale bar corresponds to a size of two kilobase pairs (Kbp). Gene cluster size is 28.207 Kbp. Abbreviations: cobP/cobU: adenosylcobinamide kinase/adenosylcobinamide-phosphate guanylyltransferase; cobA/btuR: cob(I)alamin adenosyltransferase; cobQ/cbiP adenosylcobyric acid synthase; cbiB/cobD: adenosylcobinamide-phosphate synthase; cobC1/cobC cobalamin biosynthetic protein CobC; cobC/phpB: alpha-ribazole phosphatase; cobS/cobV adenosylcobinamide-GDP ribazoletransferase; cobU/cobT: nicotinate-nucleotide--dimethylbenzimidazole phosphoribosyltransferase; bluB: 5 6-dimethylbenzimidazole synthase; cobW: cobalamin biosynthesis protein CobW; cobN: cobaltochelatase CobN; cobH/cbiC: precorrin-8X/cobalt-precorrin-8 methylmutase; cobI/cbiL: precorrin-2/cobalt-factor-2 C20-methyltransferase; cobJ/cbiH: precorrin-3B C17-methyltransferase ; cobK/cbiJ :precorrin-6A/cobalt-precorrin-6A reductase ; cobL/cbiET: precorrin-6Y C5 15-methyltransferase (decarboxylating) ; cbiG: cobalt-precorrin 5A hydrolase ; cobM/cbiF: precorrin-4/cobalt-precorrin-4 C11-methyltransferase; cbiD: cobalt-precorrin-5B (C1)-methyltransferase; cobB/cbiA: cobyrinic acid a c-diamide synthase; cysG: uroporphyrin-III C-methyltransferase/precorrin-2 dehydrogenase/sirohydrochlorin ferrochelatase; CbtA: Cobalt transporter subunit A; CbtB: Cobalt transporter subunit B.

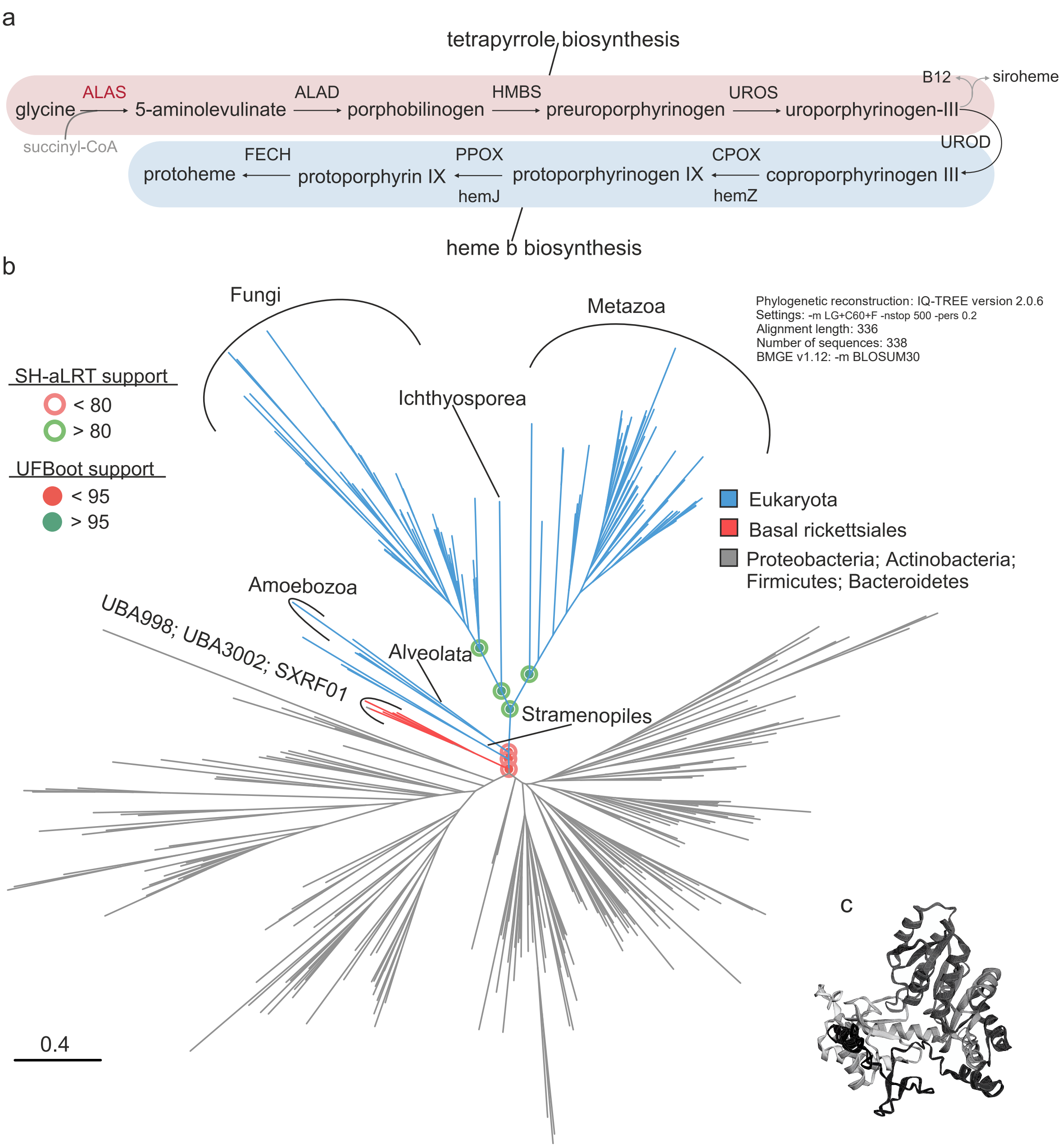


Fig. S25. **5-Aminolevulinate synthase (ALAS) gene**. a) Schematic representation of protoheme biosynthesis (heme, heme b) through C4 (Shemin pathway). The red-colored background designates the pathway segment common with adenosylcobalamin ($B_{12}$) and siroheme biosynthesis. ALAS was highlighted with red color as it represents the rate-limiting enzyme in the protoheme biosynthetic pathway. The blue-colored background denotes the protoheme-specific pathway module. Abbreviations: ALAD: porphobilinogen synthase; HMBS: hydroxymethylbilane synthase; UROS: uroporphyrinogen-III synthase; UROD: uroporphyrinogen decarboxylase; CPOX: coproporphyrinogen III oxidase; PPOX: protoporphyrinogen/coproporphyrinogen III oxidase; FECH: protoporphyrin/coproporphyrin ferrochelatase. b) Maximum-likelihood phylogeny of ALAS proteins. Eukaryotic ALAS branches (nuclear-encoded mitochondrial enzymes) are depicted in blue, while the Rickettsiales ones are in red. Non-Rickettsiales bacterial clades are colored grey (see the legend on the right side of the panel). Branch support values (UFBoot and Sh-aLRT) are indicated through colored circles (see the legend on the left side of the figure). The scale bar indicates the number of substitutions per site. c) Phyre2 predicted protein structure (100% confidence; 52% identity with ALAS from *Saccharomyces cerevisiae* S288C) of ALAS gene in basal SXRF01 Rickettsiales (RE-15aug16-6 MAG).

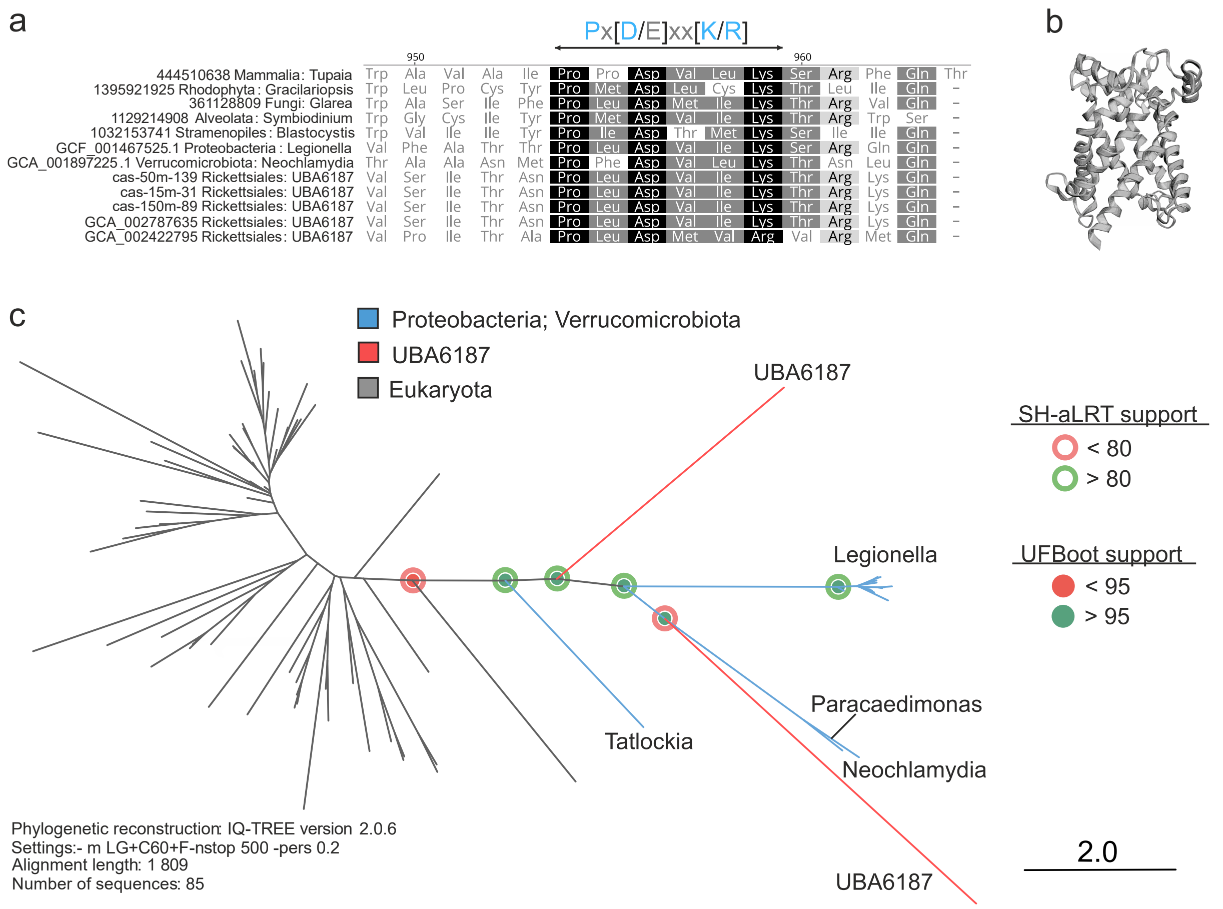


Fig. S26. **Mitochondrial carnitine/acylcarnitine transporter (CACT)**. a) Section of amino acid alignment showing the presence of the mitochondrial carrier family signature motif Px[D/E]xx[K/R] (Robinson et al., 2008). b) CACT phyre2-predicted protein structure (100% confidence; 16% identity with mitochondrial carrier protein from Mus musculus) in basal UBA6187 Rickettsiales (cas-50m-139 MAG). c) CACT maximum-likelihood phylogeny. Eukaryotic branches (nuclear-encoded mitochondrial transporters) are depicted in grey, while the Rickettsiales ones are in red. Non-Rickettsiales bacterial clades are colored in blue (see legend at the top of the panel). Branch support values (UFBoot and Sh-aLRT) are indicated through colored circles (see the legend on the right side of the figure). The scale bar indicates the number of substitutions per site.

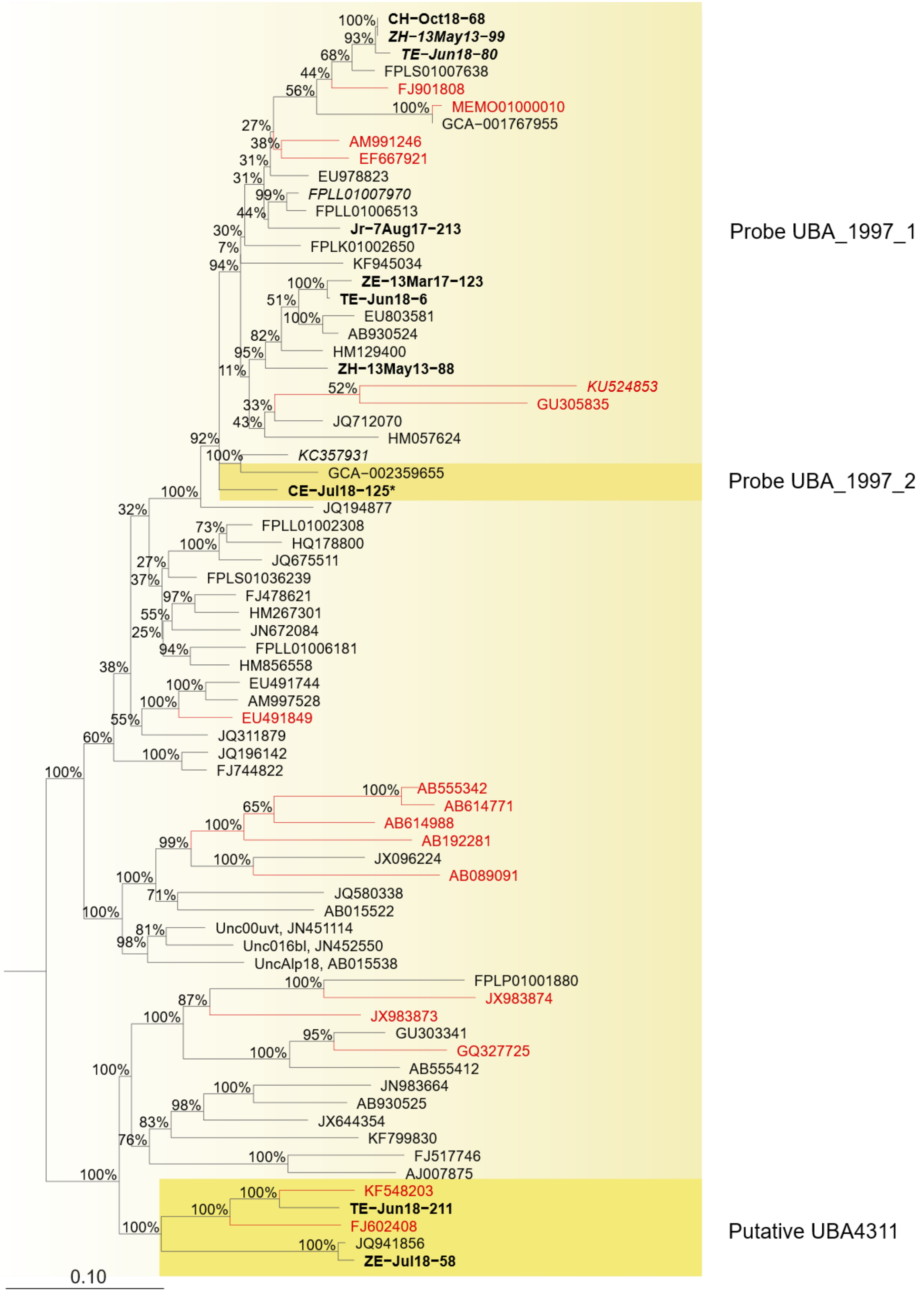


Fig. 27. **Randomized Accelerated Maximum Likelihood tree of UBA1997 16S rRNA genes**. The MAGs generated in this study are shown in bold. Sequences not targeted by CARD-FISH probes are shown in italics. *Sequences shorter than 1 000 bp that were introduced in the tree using the parsimony method in ARB software. Sequences with low

Pintail values (SILVA) are indicated in red. Branch bootstrap support values are displayed at the nods. The scale bar indicates the number of substitutions per site.

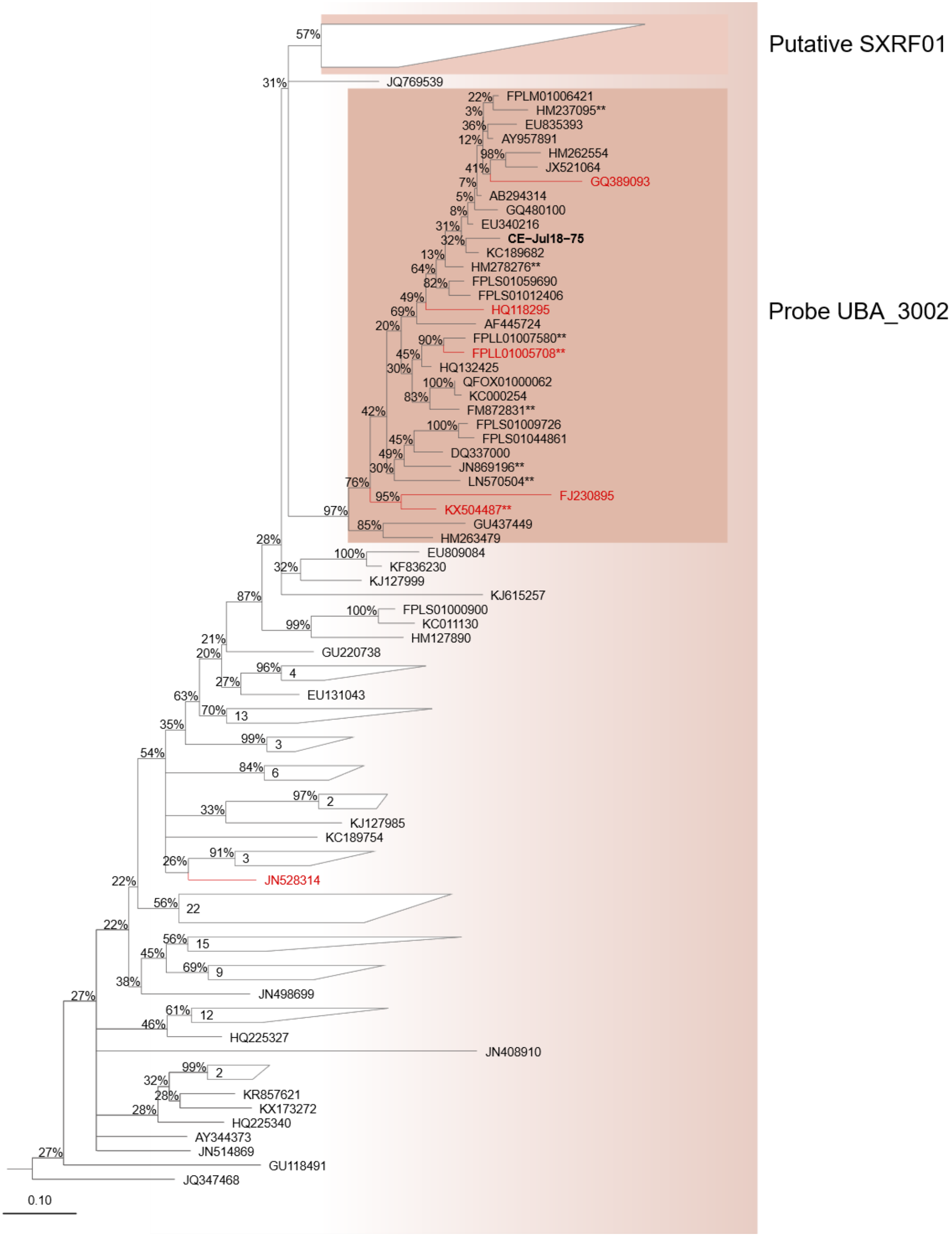


Fig. 28. **Randomized Accelerated Maximum Likelihood tree of UBA3002 and phylogenetically close 16S rRNA genes (group SM2D12, SILVA taxonomy)**. The MAGs generated in this study are shown in bold. **Sequences within the target group with mismatch to the probe. Sequences with low Pintail values (SILVA) are indicated in red. Branch bootstrap support values are displayed at the nods. Support values below 20% outside of the target group were resolved in bifurcations. Sequences outside of the target group with support values above 50% are shown as collapsed. The scale bar indicates the number of substitutions per site.

| | Metabolism | SXRF01 | UBA998 | UBA3002 | UBA8987 | 21-14-0-20-44-7 | UBA6187 | UBA1997 | UBA4311 | Midi. | Anaplas. | Arcano. | 33-17 | UBA6177 | Ricket. |
|---|---|---|---|---|---|---|---|---|---|---|---|---|---|---|---|
| Transporters | SemiSWEET transporter | | | | | | | | | | | | | | |
| | maltose/moltooligosaccharide | | | | | | | | | | | | | | |
| Catabolic | Pyruvate oxidation | | | | | | | | | | | | | | |
| | Oxidative phosphoryla-tion | | | | | | | | | | | | | | |
| | Cytochrome c oxidase cbb3 type | | | | | | | | | | | | | | |
| Anabolic | Gluconeoge-nesis | | | | | | | | | | | | | | |
| Two-compound systems | PhorR-PhorB Phosphate | | | | | | | | | | | | | | |
| | NtrB-NtrC Nitrogen | | | | | | | | | | | | | | |
| | CusR-CusS Copper | | | | | | | | | | | | | | |
| | FhaB-FhaC | | | | | | | | | | | | | | |

Table S1. **Rickettsiales genome-informed metabolic reconstructions**. Additional family-centric metabolic pathways, two-compound systems, and transporters. Rickettsiales families are separated based on their evolutionary reconstruction (from left to right): ancient (light red), transition (light yellow), and intracellular (light blue). The presence of specific pathways, two-compound systems, or transporters is depicted by dark grey (the absence is shown by the usage of light grey color). Abbreviations: Midi. = Midichloriaceae, Anaplas. = Anaplasmataceae, Ricket. = Rickettsiaceae.

| | Oxidase type | SXRF01 | UBA998 | UBA3002 | UBA8987 | 21-14-0-20-44-7 | UBA6187 | UBA1997 | UBA4311 | Midi. | Anapl. | Arcano. | 33-17 | UBA6177 | Ricket. |
|---|---|---|---|---|---|---|---|---|---|---|---|---|---|---|---|
| High affinity | bd-I | | | | | | | | | | | | | | |
| | C | | | | | | | | | | | | | | |
| Medium affinity | A1-bo3 | | | | | | | | | | | | | | |
| Low affinity | A2 | | | | | | | | | | | | | | |
| | A1 | | | | | | | | | | | | | | |

Table S2. **Cytochrome oxidase types are present in Rickettsiales genomes**. The table shows the presence (dark grey) and absence (light grey) of cytochrome oxidases classified by their oxygen affinity.

| KEGG Entry | Symbol | Name |
|---|---|---|
| K03878 | ND1 | NADH-ubiquinone oxidoreductase chain 1 |
| K03935 | NDUFS2 | NADH dehydrogenase (ubiquinone) Fe-S protein 2 |
| K03941 | NDUFS8 | NADH dehydrogenase (ubiquinone) Fe-S protein 8 |
| K02256 | COX1 | cytochrome c oxidase subunit 1 |
| K02132 | ATPeF1A, ATP5A1, ATP1 | F-type H+-transporting ATPase subunit alpha |
| K02136 | ATPeF1G, ATP5C1, ATP3 | F-type H+-transporting ATPase subunit gamma |

Table S3. **KO IDs for the six selected proteins involved in oxidative phosphorylation**.

| Probe name | Target group (SILVA taxonomy) | Sequence (5′-3′) | FA (%)/ Temp. (◦C) | Number of hits in the group (Coverage, %) | Hits in SILVA 138.1 | Hits outside the group |
|---|---|---|---|---|---|---|
| UBA_3002 | Subgroup of SM2D12 | CAAACTGGACAT GTCAAGGC | 20/35 | 24/32 (75) | 23 | 1 (AY957887, Blastomonas, Pintail value 0) |
| UBA_3002_C | competitor | CATACTGGACAT GTCAAGGG | | | | |
| UBA_1997_1<br>UBA_1997_2 | Subgroup of Mitochondria | GGTAGAAACAG CTTCCATGGC<br>GGTAGAAACGA CTTCCATGGC | 50/35<br>50/35 | 21 23/23 (100)<br>2 | 15<br>0 | 0<br>0 |
| UBA_1997_C | competitor | GGTAGAACCAA CTTCCATGGC | | | | |

Table S4. **CARD-FISH probes and competitors designed in this study**.

FA - formamide concentration used in hybridization buffer. Temp. - temperature of hybridization. Probes UBA_1997_1 and UBA_1997_2 were used together for the hybridization of the entire group.
